# Neuronal pentraxin Nptx2 regulates complement activity in the brain

**DOI:** 10.1101/2022.09.22.509106

**Authors:** Jiechao Zhou, Sarah D. Wade, David Graykowski, Mei-Fang Xiao, Binhui Zhao, Lucia AA Giannini, Jesse E. Hanson, John C van Swieten, Morgan Sheng, Paul F. Worley, Borislav Dejanovic

## Abstract

Complement overactivation mediates microglial synapse elimination in neurological diseases like Alzheimer’s disease and frontotemporal dementia (FTD), but how complement activity is regulated in the brain remains largely unknown. We identified that the secreted neuronal pentraxin Nptx2 binds complement C1q and thereby regulates its activity in the brain. Nptx2-deficient mice show increased complement activity and C1q-dependent microglial synapse engulfment and loss of excitatory synapses. In a neuroinflammation culture model and in aged TauP301S mice, AAV-mediated neuronal overexpression of Nptx2 was sufficient to restrain complement activity and ameliorate microglia-mediated synapse loss. Analysis of human CSF samples from a genetic FTD cohort revealed significantly reduced levels of Nptx2 and Nptx2-C1q protein complexes in symptomatic patients, which correlated with elevated C1q and activated C3. Together, these results show that Nptx2 regulates complement activity and microglial synapse elimination in the healthy and diseased brain and that diminished Nptx2 levels might exacerbate complement-mediated neurodegeneration in FTD patients.

## Introduction

A tight regulation of synapse formation, removal and maintenance of appropriate connections is important for the assembly of neuronal circuits in the mature brain. Aberrant synapse function and/or elimination is a key pathomechanism in neuropsychiatric and neurodegenerative disease like schizophrenia and Alzheimer’s disease (AD) (Neniskyte and Gross, 2017; Volk et al., 2014; Wilton et al., 2019). Innate immune molecules, including the classical complement pathway (CCP), play a role in synapse removal during developmental circuit refinement and in pathological conditions (Schafer et al., 2012; Stevens et al., 2007). C1q, the initiating factor of the CCP, binds to synapses and, upon activation of the downstream cascade, leads to engulfment of the complement-opsonized synapses by microglia (Stevens et al., 2007). This process is overactivated in several CNS diseases including AD, virus-induced synapse loss, frontotemporal dementia (FTD), and perhaps schizophrenia, leading to synapse loss and subsequent cognitive decline (Bohlen et al., 2019; Comer et al., 2020; Dejanovic et al., 2018; Hong et al., 2016; Vasek et al., 2016; Wu et al., 2019). Blocking C1q binding to neurons is sufficient to attenuate synapse loss in mouse models of AD (Dejanovic et al., 2018; Hong et al., 2016; Wilton et al., 2021).

Despite the pivotal role of C1q in neurodegeneration and synapse loss, it is not well understood how C1q and downstream complement activity are regulated in the brain. Recently, the secreted neuronal sushi-domain protein SRPX2 has been shown to bind C1q and block complement-mediated synapse elimination during development of the visual system and somatosensory cortex (Cong et al., 2020). However, loss of SPRX2 had no effect on complement activity and synapse density after the developmental period (Cong et al., 2020), suggesting that SPRX2 is not involved in complement regulation in the adult brain. Interestingly, APOE, a major risk factor for AD, has been shown to interact with C1q and thereby modulate CCP activity (Yin et al., 2019).

In the periphery, a number of proteins have been described to bind C1q and other complement factors and regulate complement activation. Members of the pentraxin family directly bind C1q and thereby regulate peripheral complement activity (Nauta et al., 2003). Pentraxins are characterized by their C-terminal pentraxin domain and sequence homology among family members. In the CNS, three pentraxins are primarily expressed by excitatory neurons – neuronal pentraxin 1 and 2 (Nptx1 and Nptx2), and the neuronal pentraxin receptor (Nptxr). NPTXs form disulfide-linked hetero-oligomers that remain tethered to the plasma membrane by the transmembrane domain of Nptxr (Xu et al., 2003). NPTX oligomers bind and promote the clustering of postsynaptic AMPA type glutamate receptor (AMPA-R), thereby contributing to different forms of synaptic plasticity (Chang et al., 2010; Cho et al., 2008; Lee et al., 2016; O’Brien et al., 1999; Pelkey et al., 2016; Sia et al., 2007). Moreover, Nptxr-containing complexes can be shed from the synapse upon enzymatic cleavage of Nptxr (Cho et al., 2008). In vivo imaging of Nptx2 indicates that the Nptx1/2/r complex is exocytosed at excitatory synapses on parvalbumin interneurons (PV-IN) during periods of behavioral activity and as much as 40% is later shed on a diurnal and circadian schedule during sleep, presumably in the process of circuit refinement (Xiao et al., 2021). The shedding process can be monitored in human subjects by measuring NPTX2 in CSF and is disrupted by sleep deprivation in both mouse and human subjects highlighting its dynamic physiological regulation (Xiao et al., 2021).

Nptx2 is expressed as an immediate early gene by excitatory neurons and is particularly important for activity-dependent strengthening of GluA4 AMPA-R containing excitatory synapses on PV-IN (Chang et al., 2010; Sia et al., 2007; Xiao et al., 2017). Ocular dominance plasticity requires NPTX2 (Gu et al., 2013) and monocular deprivation results in rapid shedding of Nptx2 expressed by pyramidal neurons at excitatory synapses on PV-IN in deprived cortex with co-incident disconnection of excitatory synapses coupling pyramidal neurons with PV-IN (Severin et al., 2021). In an AD amyloidosis mouse model, Nptx2 amplifies the effect of amyloid-beta (Aß) to reduce inhibitory circuit function required for gamma rhythmicity (Xiao et al., 2017). Nptx2 in CSF serves as a biomarker in genetic FTD (Ende et al., 2020), AD (Galasko et al., 2019; Xiao et al., 2017), Down syndrome (Belbin et al., 2020), Dementia with Lewy Bodies (Steenoven et al., 2020), schizophrenia (Xiao et al., 2021) and normal aging (Soldan et al., 2019), and consistently correlates with cognitive measures. The reduction of Nptx2 in CSF may reflect inhibitory circuit dysfunction with associated excitatory synapse damage or loss, a pathomechanism common between neurodegenerative and neuropsychiatric diseases (Styr and Slutsky, 2018; Xiao et al., 2017, 2021).

Here, we examined the hypothesis that synaptic NPTX proteins might bind C1q and regulate the CCP in the brain. We provide evidence that Nptx2 binds C1q and thereby attenuates complement activity in the adult mouse brain. Deletion of Nptx2 leads to increased activation of the CCP and microglia-mediated elimination of excitatory synapses. These consequences of Nptx2^KO^ are prevented by genetic deletion of C1q or C1q blocking antibody, consistent with the notion that the natural function of Nptx2 is to inhibit C1q and classical complement activation in brain. Nptx2 also inhibits the cytotoxic action of activated microglia in a co-culture system. Moreover, in TauP301S mice, a FTD and AD model characterized by neuroinflammation and complement-dependent neuronal damage (Dejanovic et al., 2018; Wu et al., 2019), AAV-mediated overexpression of Nptx2 decreased complement activation and associated synapse loss. Lastly, we show that Nptx2 interacts with C1q in human CSF and that the fraction of Nptx2-bound C1q is decreased in symptomatic carriers of genetic FTD, while complement activity is increased compared to healthy non-carriers and presymptomatic carriers.

## Results

### Neuronal pentraxins bind C1q and inhibit CCP activity in vitro

C1q binds peripheral pentraxins with high affinity (Inforzato et al., 2006). To determine if C1q interacts with neuronal pentraxins, we immobilized purified NPTXs, or IgM as a positive control, onto microtiter plate wells and measured C1q binding. C1q bound to all three NPTXs (Fig. 1A). Binding to immobilized Nptxr was protein-concentration dependent and saturable (Fig. S1A). The interaction was C1q-specific as NPTXs did not bind to immobilized complement C2, C3 or C4 (Fig. 1B). To visualize the C1q-Nptx2 complex, we analyzed purified Nptx2 and C1q by negative staining electron microscopy. As expected, C1q showed a characteristic radial spoke structure with six globular heads connected by collagen-like stalks, while Nptx2 formed a hexameric ring ∼10nm across (Fig. 1C). When incubated together, the resulting C1q-Nptx2 complex showed Nptx2 bound to the core of C1q molecule in-between the globular heads (Fig. 1C). Peripheral pentraxins bind C1q with their pentraxin domain (Inforzato et al., 2006). To test if C1q binds to the pentraxin domain of NPTXs, we expressed and purified Nptx2 pentraxin domain (X2-PD) and immobilized it to Cyanogen bromide-activated (CNBr) Sepharose beads. X2-PD coated beads, but not control beads, pulled down C1q (Fig. S1B). To independently confirm the C1q-NPTX interaction, we performed cell-surface binding assays with a stable tetracycline-inducible CHO cell-line that co-expresses Nptxr and Nptx2 (Nptxr/x2). Upon addition of C1q to fresh cell medium, essentially all CHO cells with tetracycline-induced Nptxr/x2 expression were decorated with C1q, while C1q did not bind to the CHO cell-lines in absence of tetracycline (and without detectable expression of Nptxr and Nptx2) (Fig. 1D). To estimate the C1q-Nptxr/x2 binding affinity, we incubated Nptxr/x2-expressing CHO cells with increasing concentrations of C1q, fixed the cells and quantified the amount of bound C1q on the surface using an anti-C1q antibody. We measured a saturable interaction with a K_D_ ∼2 nM for C1q and Nprtx/x2 (Fig. 1E).

**Figure 1:**
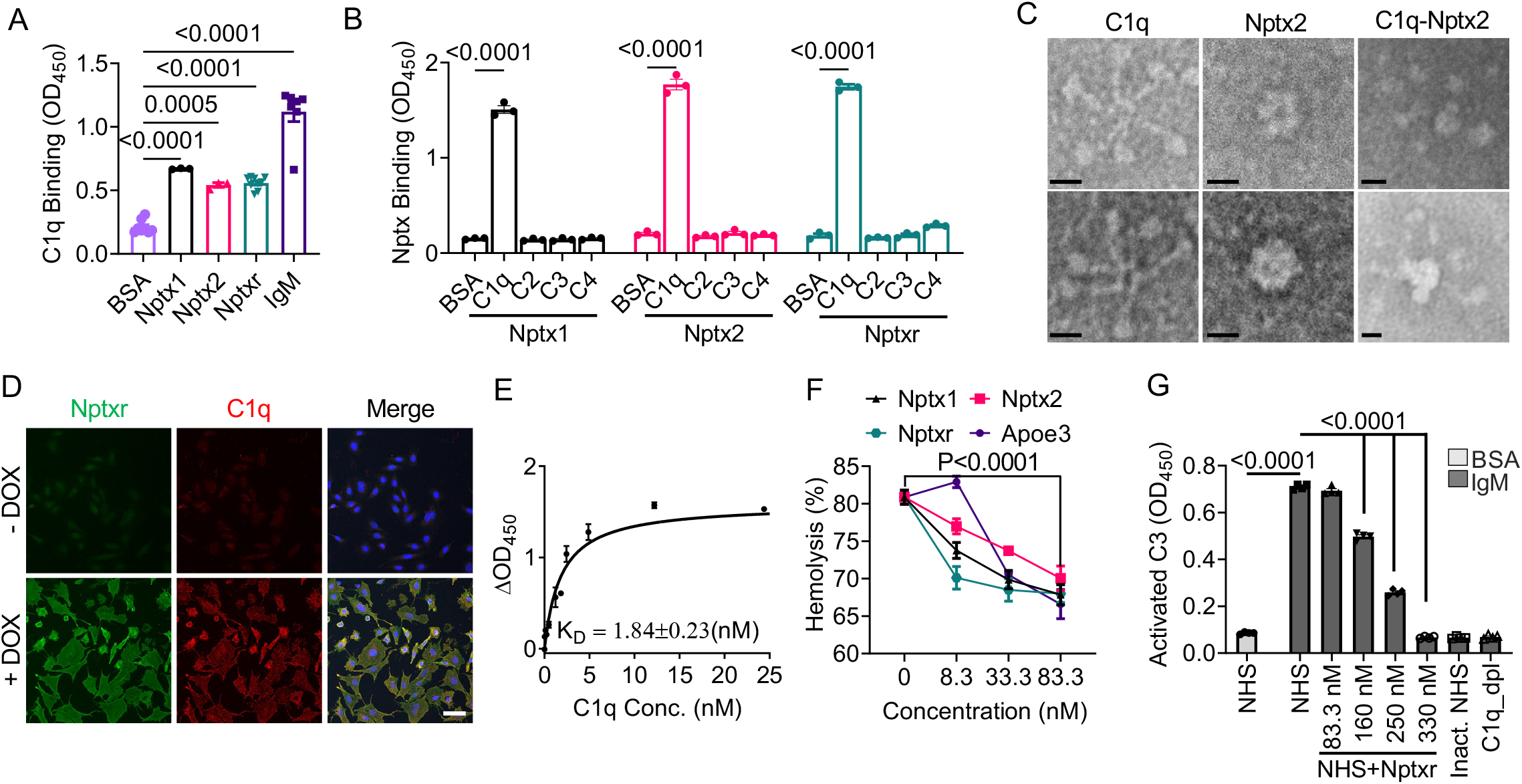
Neuronal pentraxins bind C1q and inhibit CCP activity *in vitro*. A) Microtiter wells were coated with BSA, IgM or neuronal pentraxins (Nptx1, Nptx2, Nptxr) as indicated and incubated with C1q. Binding of C1q to the wells was detected using an anti-C1q antibody. p values were determined by one-way ANOVA. B) Microtiter wells were coated with BSA or complement proteins (C1q, C2, C3, C4) as indicated. Binding of his-tagged neuronal pentraxins was monitored using an anti-His antibody. p values were determined by Two-way ANOVA. C) Representative images of negative staining electron microscopy of purified C1q, Nptx2, and C1q-Nptx2 complex. Scale bar: 10 nm. D) C1q cell-surface binding assays with a stable tetracycline-inducible CHO cell-line that co-expresses Nptxr and Nptx2. C1q proteins were added to the fresh culture medium, and bound C1q was visualized by immunofluorescence labeling (red). Expression of Nptxr was detected with a Nptxr-specific antibody (green). Scale bar: 50 µm. E) Cell-surface binding to Nptxr/x2-expressing CHO cells with C1q as ligand. Stable inducible CHO cells expressing Nptxr/x2 were incubated with C1q at indicated concentrations. Bound C1q was detected with an anti-C1q antibody. The signal shown was subtracted from the background in CHO cells without induced Nptxr expression. p value was determined by Two-way ANOVA. F) Neuronal pentraxin inhibits hemolysis of sheep erythrocytes. Purified Nptx1, Nptx2, Nptxr or ApoE3 was incubated in normal human serum (NHS), which was activated via CCP-specific GVB++ buffer, and lysis of sheep erythrocytes was analyzed by measuring released hemoglobin at 415 nm. G) Nptxr inhibits CCP and activated C3 deposition. 1% NHS supplemented with recombinant Nptxr was added to IgM-coated microtiter plates and activated C3 deposition was measured using specific antibodies. Heat-inactivated NHS and C1q-depleated human serum were used as negative control. p values were determined by one-way ANOVA. Data are presented as mean ± SEM. Each dot represents data from individual samples.

Based on the interaction data, we tested if NPTXs modulate C1q-inducted CCP activity. In a hemolytic complement activity assay, addition of normal human serum (NHS), which contains all CCP proteins, leads to the lysis of erythrocytes (Fig. S1C). Similar to ApoE, which was previously shown to inhibit CCP (Yin et al., 2019), addition of Nptx1, Nptx2 or Nptxr decreased complement-mediated hemolysis in a dose-dependent manner (Fig. 1F). Addition of GST-tagged Nptxr pentraxin domain (XR-PD), but not GST alone, efficiently inhibited CCP-mediated hemolysis in a dose-dependent manner (Fig. S1D). Furthermore, in a CCP-mediated *E. coli* killing assay, bacteria cells remained viable in presence of XR-PD in a dose-dependent manner (Fig. S1E). Lastly, we used a plate-based assay in which we immobilized IgM onto microtiter plate wells. C1q binds with high affinity to aggregated antibodies resulting in the activation of the CCP; addition of NHS with CCP-activating buffer showed potent C1q-dependent deposition of cleaved, i.e. activated, C3 in IgM coated wells, reflecting CCP activation. Addition of Nptxr inhibited CCP activity in a dose-dependent manner (Fig. 1G). Together, these results show that all three NPTXs can bind C1q and inhibit the CCP, likely by sequestering and blocking C1q.

### Nptx2 deletion increases CCP activity and leads to loss of excitatory synapses in vivo

In AD, Nptx2 is decreased in postmortem brain and in patient CSF where its levels correlate with disease status, cognitive performance and disease progression (Xiao et al., 2017). Nptx2 levels are also reduced in CSF from schizophrenia patients, and Nptx2 knockout (Nptx2^KO^) mice show schizophrenia-related behavioral deficits (Xiao et al., 2021). Given the identification of Nptx2 as a regulator of complement, we next examined if Nptx2^KO^ mouse brains exhibited dysregulated complement. We first immunostained C1q in Nptx2^KO^ mouse brains; C1q immunofluorescence intensity was similar in Nptx2^KO^ compared to WT brains (Fig. 2A), which we further validated by C1q ELISA (Fig. 2B). We next examined C4 which is downstream of C1q in the CCP. While full-length C4 levels were also unchanged in Nptx2^KO^ brains (Fig. 2C), cleaved C4b levels were significantly increased in Nptx2^KO^ brains (Fig. 2D), which is consistent with increased processing of C4 resulting from increased CCP activity in NPTX2^KO^ brains.

**Figure 2:**
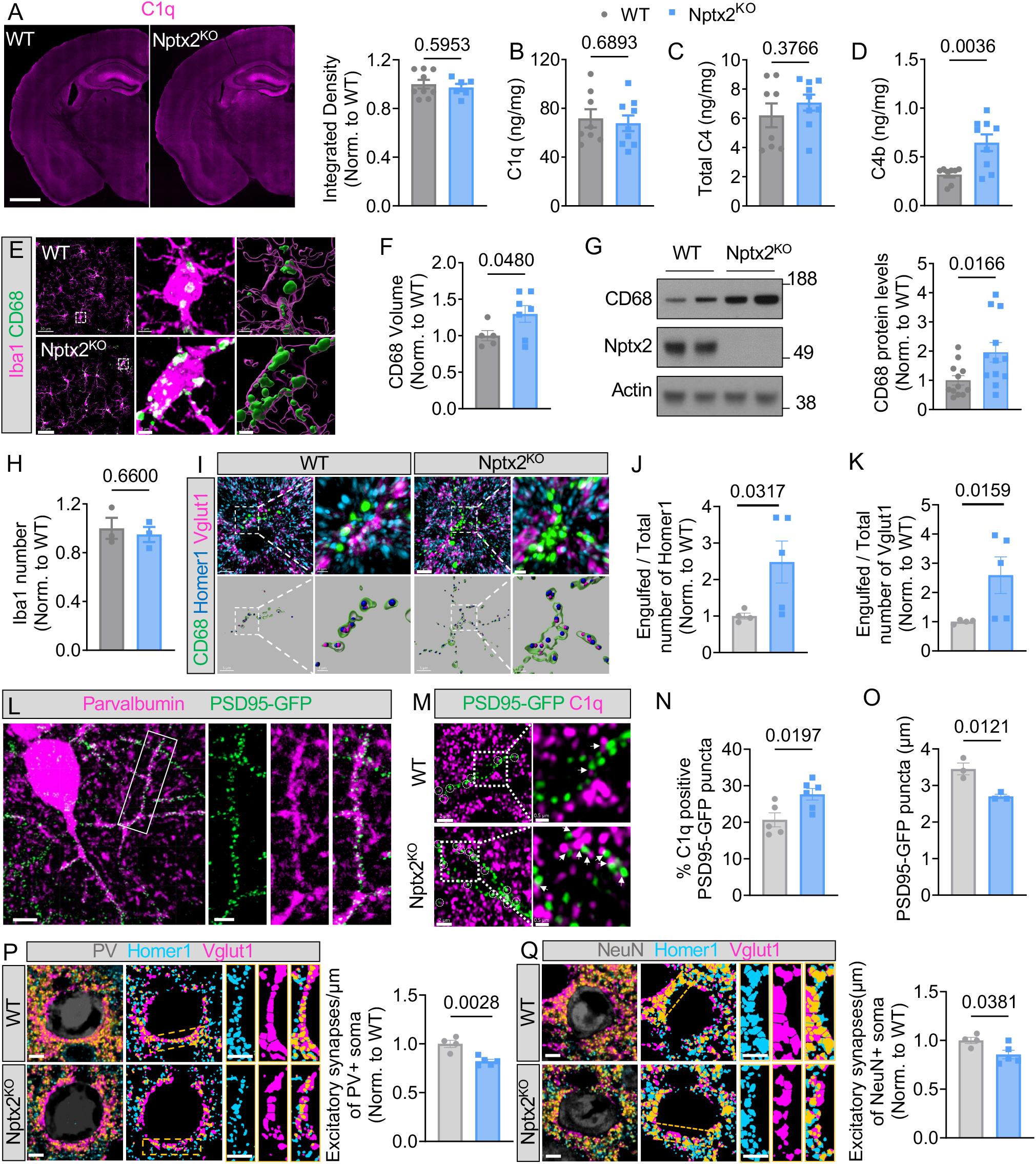
Nptx2 regulates complement activity *in vivo*. A). Representative images of C1q immunostaining in WT and Nptx2^KO^ mice brains. Scare bar is 1000 µm; Right panel: Quantification of C1q immunofluorescence in WT and Nptx2^KO^ brains. p values were determined by unpair *t* test. n = 9 for WT; n = 6 for Nptx2^KO^. B-D) C1q, C4, and C4b concentration in WT and Nptx2^KO^ cortex was measured by ELISA. p values were determined by unpair *t* test. n = 8 for WT; n = 9 for Nptx2^KO^. E) Representative confocal images of CD68 (green) and Iba1 (magenta) immunostaining in WT and Nptx2^KO^ cortex. Scale bar is 30 μm; inset, 2 µm. F) Quantification of CD68 volume in WT and Nptx2^KO^ cortex. p values were determined by unpair *t* test. n = 5 for WT; n = 7 for Nptx2^KO^. G) Representative Western blots and quantification of CD68 levels in hippocampi from WT and Nptx2^KO^ cortex. p values were determined by unpair *t* test. n = 12 for WT; n = 12 for Nptx2^KO^. H) Number of Iba1+ microglia in WT and Nptx2^KO^ whole brains. p values were determined by unpair *t* test. n = 3 for WT; n = 3 for Nptx2^KO^. I-K) Representative confocal images and 3D surface rendering of immunostained CD68 (green), Vglut1 (magenta) and Homer1 (blue) in WT and Nptx2^KO^ cortex. Scale bars are 5 µm; inset, 1 µm. Quantification of (J) Homer1 and (K) Vglut1 puncta inside CD68+ lysosomes. p values were determined by unpair *t* test. n = 4 for WT; n = 5 for Nptx2^KO^. L) Representative confocal images of PSD95-GFP and immunostained parvalbumin in PSD95-GFP^f/f^:PV-Cre WT cortex. Scale bars is 10 μm; inset, 4 μm. M) Representative confocal images of immunostained C1q (magenta) and PSD95-GFP (green) in PSD95-GFP^f/f^:PV-Cre:WT and Nptx2^KO^ hippocampus. White circles indicate C1q colocalized with PSD95-GFP. Scale bar is 2 μm; inset, 0.5 μm. N) Percentage of C1q+ PSD95-GFP puncta in WT and Nptx2^KO^ hippocampus. p values were determined by unpair *t* test. n = 5 for WT; n = 6 for Nptx2^KO^. O) PSD95-GFP puncta density in PSD95-GFP^f/f^:PV-Cre:WT and Nptx2^KO^ hippocampus. p values were determined by unpair *t* test. n = 3 for WT; n = 3 for Nptx2^KO^. P) Representative confocal images of immunostained PV (gray), Homer1 (blue), Vglut1 (magenta), and corresponding masks. Bar graph shows excitatory synapse density around somas of PV+ cells. p values were determined by unpair *t* test. n = 4 for WT; n = 5 for Nptx2^KO^. Scale bars are 5 µm. Q) Representative confocal images and mask of immunostained NeuN (gray), Homer1 (blue), Vglut1 (magenta) in WT and Nptx2^KO^ cortex. Scale bars is 5 µm. Bar graph shows normalized number of excitatory synapses around NeuN+ soma. p values were determined by unpair *t* test. n = 4 for WT; n = 5 for Nptx2^KO^. Scale bars are 5 µm. Data are presented as mean ± SEM. Each dot represents data from one mouse.

Microglia are effectors of complement-mediated synapse pruning. Given the increased complement activity, we wondered if microglia are activated in Nptx2^KO^ brains. By IHC staining and immunoblotting, we found that levels of the lysosomal marker CD68, a marker of microglial activation, were significantly increased in the cortex from Nptx2^KO^ mice (Fig. 2E-G). The number of microglia was unchanged in the cortex of Nptx2^KO^ brains (Fig. 2H). By IHC, we confirmed that Nptx2 is expressed by excitatory neurons (Fig. S2A, B) and detected secreted Nptx2 puncta in the vicinity of excitatory synapse marker proteins Vglut1 and Synaptophysin, but not the inhibitory synapse marker protein Gephyrin (Fig. S2C, D). Thus, we wondered if loss of Nptx2 might render synapses more vulnerable to complement-dependent synapse loss through microglial phagocytosis. By co-immunostaining of excitatory postsynaptic marker Homer1 and presynaptic marker Vglut1 with microglial CD68^+^ lysosomes (Fig. 2I), we found significantly more Homer1 and Vglut1 immunoreactivity inside CD68^+^ lysosomes in the cortex from Nptx2^KO^ mice compared to WT brains (Fig. 2 J, K), suggesting that microglial phagocytosis of excitatory synapses is increased in Nptx2^KO^ brains. Given that Nptx2 concentration is particularly high on dendrites of PV-INs (Chang et al., 2010), we hypothesized that excitatory synapses on PV-INs might be especially vulnerable to complement-mediated pruning in Nptx2^KO^ mice. Using mice that selectively express GFP-tagged PSD95 in PV-INs (Fig. 2L), we found a significant increase in C1q-labeled PSD95-GFP puncta in Nptx2^KO^ mice (Fig. 2M, N). The increase in C1q+ synapses correlated with decreased PSD95-GFP density along dendrites in Nptx2^KO^ brains (Fig. 2O). We speculated that microglia eliminate the excitatory synapses on PV-INs; however, we failed to identify GFP fluorescence in microglial lysosomes, possibly because GFP fluorescence is pH sensitive. Instead, using a PV-Cre reporter mouse that expresses the pH-stable tdTomato selectively in PV-INs, we found significantly more tdTomato^+^ structures within CD68+ microglial lysosomes in NPTX2^KO^ vs WT brains (Fig. S2E). To validate the loss of excitatory synapses on PV-INs, we measured the density of co-localized postsynaptic Homer1 and presynaptic Vglut1 puncta around PV somas. Excitatory synapse density around PV somas was significantly decreased in Nptx2^KO^ brains by ∼20% (Fig. 2P). PV cell density was unaffected, suggesting that microglia remove synapses (and possibly other structures) from PV-INs without inducing PV neuronal loss in Nptx2^KO^ brains (Fig. S2F). To test if excitatory synapses are more globally affected in Nptx2^KO^ brains, we measured excitatory synapse density around NeuN+ somas (a pan-neuronal marker). Intriguingly, excitatory synapse density was significantly decreased by ∼15% around NeuN+ somas (Fig. 2Q), suggesting that loss of Nptx2 leads to loss of excitatory synapses from additional neuronal subtypes beyond PV-INs.

Microglia have been shown to prune inhibitory synapses under physiological and pathological conditions through a complement-dependent mechanism (Favuzzi et al., 2021; Lui et al., 2016). To determine if loss of Nptx2 has an impact on inhibitory synapse density, we immunostained the inhibitory postsynaptic marker Gephyrin and presynaptic marker GAD67 and measured inhibitory synapse density around PV+ and NeuN+ somas (Fig. S3A, C). Inhibitory synapse density was not significantly changed in Nptx2^KO^ vs WT brains (Fig. S3B, D). The amount of Gephyrin puncta inside microglial lysosomes was comparable between WT and Nptx2^KO^ brains, whereas GAD67 content was slightly elevated in Nptx2^KO^ microglial lysosomes (Fig. S3E, F).

Taken together, our results show that microglia remove excitatory synapses in the Nptx2^KO^ cortex, whereas inhibitory synapses are largely unaffected by Nptx2 deficiency.

### Complement inhibition rescues synapse loss in Nptx2^KO^ brains

To test if synapse loss in Nptx2^KO^ brains is complement-dependent, we compared animals that were deficient in C1q or Nptx2 or in both (Nptx2^KO^;C1q^KO^ double knockout mice). Given that the volume of microglial CD68+ lysosomes was significantly increased in Nptx2^KO^ brains, we first analyzed if this microglial phenotype is C1q dependent. CD68 volume in C1q^KO^ brains was unchanged compared to WT mice, but C1q deletion prevented the CD68 increase in Nptx2^KO^ brains (Fig. S4A). Concordantly, microglial excitatory synapse engulfment, as measured by density of Homer1 and Vglut1 puncta inside CD68+ lysosomes, was unchanged in C1q^KO^ compared to WT brains, but was significantly decreased in Nptx2^KO^;C1q^KO^ vs Nptx2^KO^ brains (Fig. 3A-C). To measure if C1q deletion in Nptx2^KO^ brains resulted in increased synapse density, we measured excitatory synapse density around NeuN+ and PV+ somas. At 10 weeks age, C1q^KO^ mice had a comparable synapse density to WT mice around NeuN+ and PV+ somas, suggesting that C1q is not essential for synapse pruning in the adult brain (Fig. 3D-G). Importantly, the decreased synapse density seen in Nptx2^KO^ mice was rescued by C1q deletion, as Nptx2^KO^;C1q^KO^ mice had a similar synapse density to WT mice (Fig. 3D-G). Since in all these IHC readouts C1q^KO^ and Nptx2^KO^;C1q^KO^ brains were not statistically different, the data imply that C1q is required in Nptx2^KO^ to induce microglia activation and excitatory synapse elimination.

**Figure 3:**
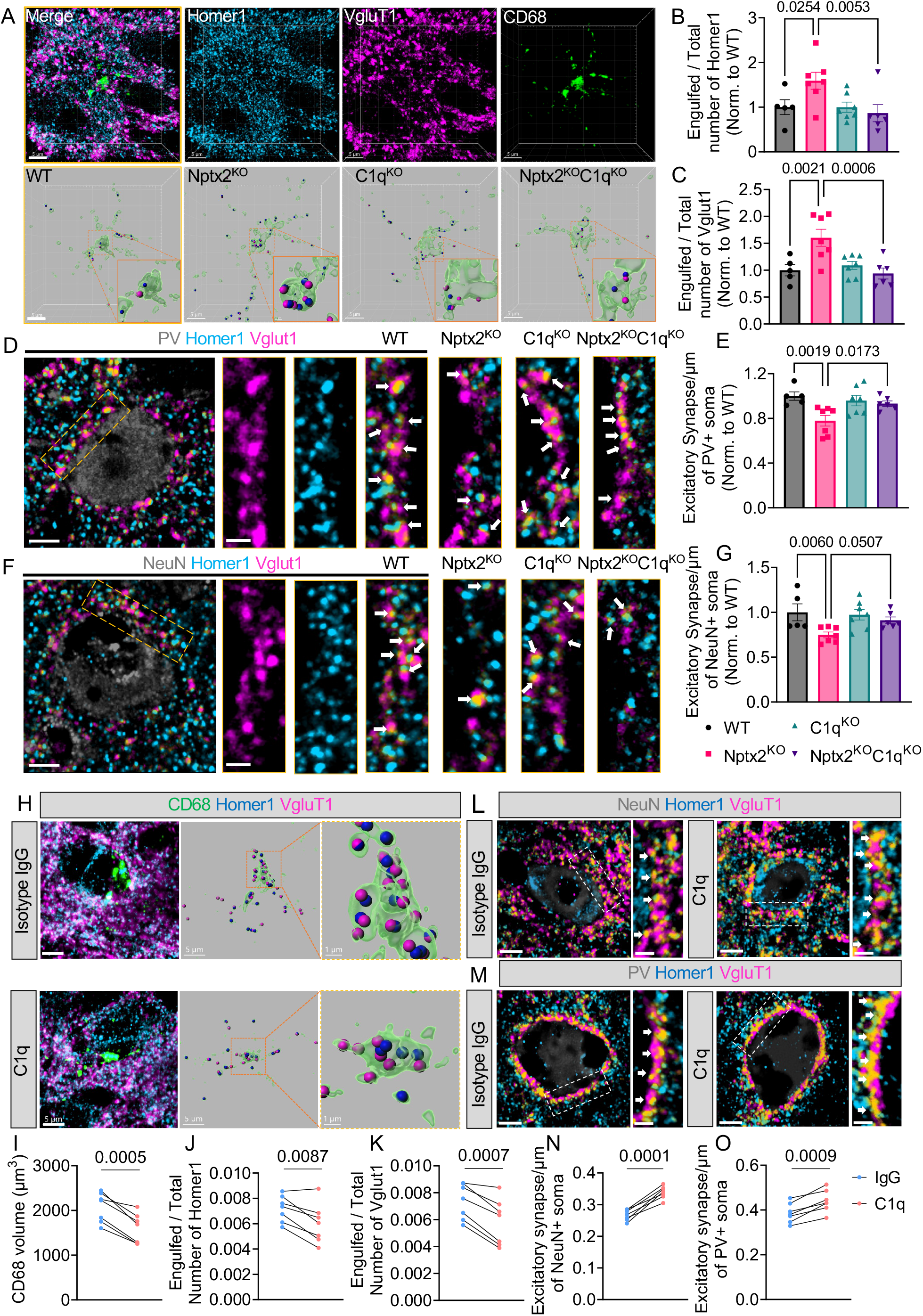
Complement inhibition rescues synapse loss in Nptx2^KO^ brains. A) Representative confocal image and 3D surface reconstruction of Homer1 (blue), Vglut1 (magenta) and CD68 (green) in the mouse cortex. Scale bars are 5 µm. B, C) Relative amount of Homer1 and Vglut1 puncta within CD68+ microglial lysosomes. p values were determined by one way ANOVA. n = 5 for WT; n = 7 for Nptx2^KO^; n = 7 for C1q^KO^; n = 6 for Nptx2^KO^;C1q^KO^. D, E) Representative confocal images of Homer1, Vglut1 and PV. Bar graph shows excitatory synapse density around somas of PV+ cells in the mouse cortex. p values were determined by one way ANOVA. n = 5 for WT; n = 7 for Nptx2^KO^; n = 7 for C1q^KO^; n = 6 for Nptx2^KO^;C1q^KO^. Scale bar is 5 µm; inset: 2 µm. F, G) Representative confocal images and quantification of excitatory synapse density around somas of NeuN+ cells in the cortex. White arrows indicate synapses, which were identified by colocalization of Homer1 and Vglut1 puncta. p values were determined by one way ANOVA. n = 5 for WT; n = 7 for Nptx2^KO^; n = 7 for C1q^KO^; n = 6 for Nptx2^KO^;C1q^KO^. Scale bar is 5 µm; inset. 2 µm. H) Representative confocal images and 3D surface reconstruction of CD68 (green), Vglut1 (magenta) and Homer1 (blue) in control IgG and C1q-blocking antibody injected Nptx2^KO^ cortex. Scale bar is 5µm; inset, 1µm. I) Volume of CD68+ microglial lysosomes in the cortex of Nptx2^KO^ mice injected with control IgG and C1q-blocking antibody. p value was determined by paired *t* test. n = 7. J, K) Relative amount of Homer1 and Vglut1 puncta within CD68+ microglial lysosomes in the cortex of Nptx2^KO^ mice injected with isotype IgG antibody or C1q-blocking antibody. p values were determined by paired *t* test. n = 7. L) Representative confocal images of immunostained NeuN (grey), Homer1 (blue) and Vglut1 (magenta) in the cortex of Nptx2^KO^ mice injected with control IgG or C1q-blocking antibody. White arrows indicate synapses, which were identified by colocalization of Homer1 and Vglut1. p value was determined by paired *t* test. n = 7; Scale bar is 5 µm; inset: 2 µm. M) Representative confocal images of immunostained PV (grey), Homer1 (blue) and Vglut1 (magenta) in the cortex of Nptx2^KO^ mice injected with control IgG or C1q-blocking antibody. White arrows indicate synapses, which were identified by colocalized Homer1 and Vglut1 puncta. Scale bar is 5 µm; inset, 2 µm. N) Excitatory synapse density around somas of NeuN+ cells. p value was determined by paired *t* test. n = 7; O) Excitatory synapse density around somas of PV+ cells. p value was determined by paired *t* test. n = 7; Data are presented as mean ± SEM. Each dot represents data from one mouse.

We showed previously that C1q-blocking antibodies rescued synapse loss in neuron-microglia cultures and in P301S mice (Dejanovic et al., 2018). To examine if acute inhibition of C1q is sufficient to reduce microglial synapse elimination and rescue synapse density, we stereotactically injected C1q-blocking antibodies, that potently inhibit CCP activity in vitro (Fig. S4B), or isotype control IgGs into opposite cortical hemispheres of Nptx2^KO^ mice and analyzed the brains 6 days post injection (Fig. S4C,D). While the control IgG was diffusively distributed in the hippocampus and cortex, the anti-C1q antibody was strongly enriched in the hippocampus and showed a punctate staining that is characteristic for C1q distribution in the brain (Fig. S4E) (Dejanovic et al., 2018). By IHC, we found a significant decrease in CD68 volume in the C1q-blocking antibody injected cortex compared to the contralateral control IgG-injected cortex (Fig. 3H, I). Furthermore, microglial engulfment of the excitatory synapse marker proteins Homer1 and Vglut1 was significantly decreased in C1q-blocking vs. control IgG-injected cortical hemispheres (Fig. 3H, J, K). To determine if decreased synapse engulfment upon C1q-blocking antibody injection results in increased synapse density, we measured Homer1-Vglut1 colocalized puncta around NeuN+ and PV+ somas 6 days after antibody injection. A single dose of C1q-blocking antibody was sufficient to increase synapse density around NeuN+ and PV+ somas compared to IgG-injected contralateral cortical hemispheres in all Nptx2^KO^ mice used in the experiment (Fig. 3L-O).

Together, these findings identify that Nptx2 deletion leads to C1q/CCP-dependent synapse elimination through microglial pruning in the adult brain.

### Neuronal Nptx2 overexpression limits complement-mediated neurotoxicity in a neuroinflammation neuron-microglia co-culture model

Activated microglia can mediate neurodegeneration in culture. Addition of the bacterial endotoxin lipopolysaccharide (LPS) to neuron-microglia co-cultures induces a strong proinflammatory and phagocytic microglia state (Pulido-Salgado et al., 2018), leading to dramatic loss of neurons and dendrites as measured by Map2 immunostaining (S5A, B). It has been shown previously that blocking C1q is sufficient to ameliorate neurotoxicity mediated by activated microglia (Ndoja et al., 2020). Strikingly, when we added C1q-blocking antibodies together with LPS to neuron-microglia co-cultures, neuronal loss was completely prevented (Fig. S5A, B). Thus, C1q-mediated activation of CCP is essential for LPS-induced neurotoxicity in those co-cultures. Given that Nptx2 binds C1q and blocks CCP activation, we wondered whether increased Nptx2 levels would be sufficient to prevent LPS-induced neurotoxicity in neuron-microglia co-cultures. We added recombinant Nptx2 protein (rNptx2) or BSA as a negative control together with LPS to the cultures (Fig. S5C). Interestingly, addition of rNptx2 prevented neurotoxicity induced by LPS (Fig. S5C, D). To test whether enhanced Nptx2 expression by neurons would be sufficient to ameliorate neurotoxicity, we infected neuronal cultures with adeno-associated viruses (AAVs) overexpressing V5-tagged Nptx2 or GFP as a control for 3 days prior to the addition of microglia to the cultures. AAV-V5-Nptx2 resulted in an increased expression and secretion of Nptx2 into the medium (Fig. S5E). Remarkably, overexpression of Nptx2, but not GFP, prevented neuronal killing by LPS-activated microglia (Fig. 4A, B). We speculated that secreted Nptx2 bound and sequestered extracellular C1q thereby inhibiting CCP activation. To test this hypothesis, we established a proximity ligation assay (PLA) utilizing DNA-labeled antibodies, followed by ligation of the two DNA-tails, amplification and detection by qPCR to detect and quantify levels of C1q-Nptx2 complex in the culture medium (see Methods). Indeed, AAV-mediated overexpression of Nptx2 or the addition of rNptx2 significantly increased C1q-Nptx2 complex levels compared to the respective controls in the co-culture medium (Fig. 5C, S5F). These results show that, analogous to C1q-blocking antibodies, secreted Nptx2 can bind and block C1q, thereby preventing neuronal loss in microglia-neuron cocultures by acting as a decoy receptor.

**Figure 4:**
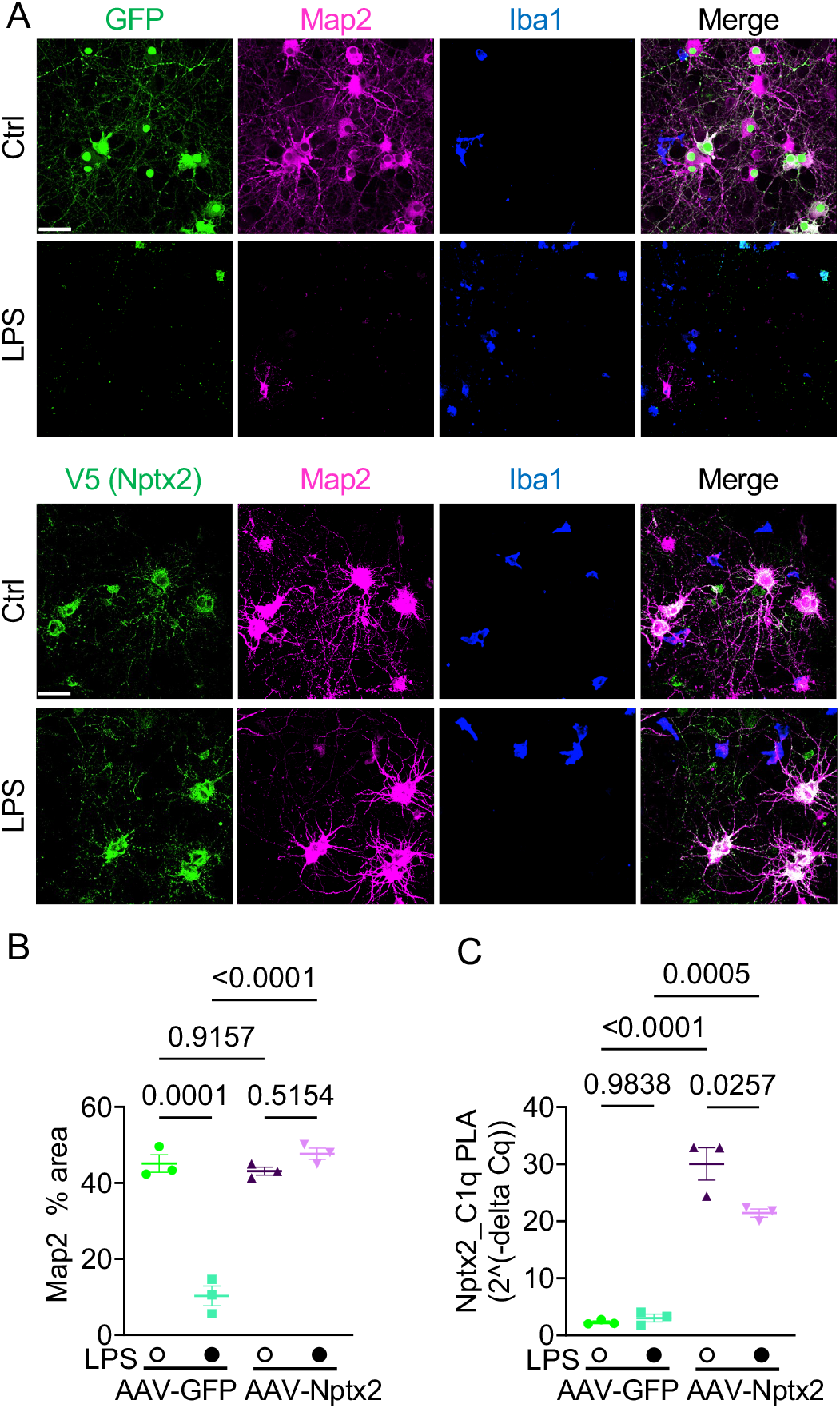
Neuronal Nptx2 overexpression limits complement-mediated neurotoxicity in a neuroinflammation neuron-microglia co-culture model. A) Representative images of immunostained Map2, Iba1 and Nptx2-V5 or GFP fluorescence in neuron-microglia co-cultures treated with LPS or vehicle for 24 hours. Neurons were infected with AAV-GFP or AAV-Nptx2-V5. Scale bar is 50 μm. B) Percentage of Map2+ area in indicated cultures. Three independent neuron-microglia co-cultures were used per condition. p values were determined by two-way ANOVA. C) Nptx2-C1q complex levels in culture medium as analyzed by proximity ligation assay (PLA, see Methods). Media from three independent neuron-microglia co-cultures were used. p values were determined by two-way ANOVA.

**Figure 5:**
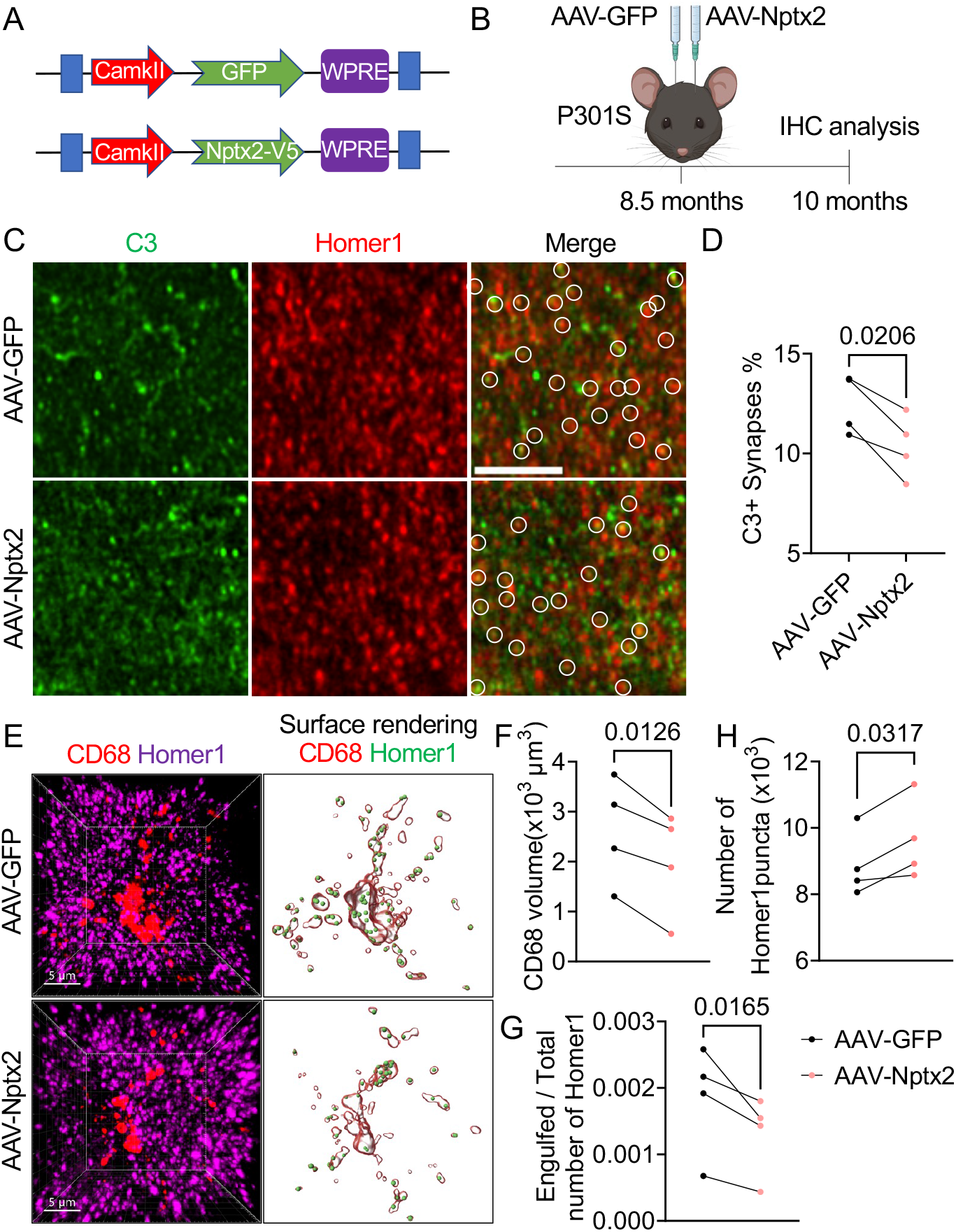
AAV-induced Nptx2 overexpression ameliorates synapse loss in P301S mice. A) Schematic representation of AAV-CamKII-GFP and AAV-CamKII-Nptx2-V5 constructs used for in vivo studies. B) Schematic illustration of the workflow and data analysis of AAV injection into brains of P301S mice. C) Representative confocal images of immunostained C3 (green) and Homer1 (red) in AAV-GFP and AAV-Nptx2 injected hippocampal regions of P301S mice. White circles indicate C3 colocalized with Homer1. Scale bar, 5 µm. D) Percentage of C3-labeled Homer1 puncta in AAV-GFP and AAV-Nptx2 injected hippocampi from P301S mice. p value was determined by paired *t* test. n = 4. E) Representative confocal images and 3D surface rendering of Homer1 and CD68+ microglial lysosomes in AAV-GFP and AAV-Nptx2 injected hippocampi of P301S mice. F) Volume of CD68+ structures in AAV-GFP and AAV-Nptx2 injected hippocampi of P301S mice. p value was determined by paired *t* test. n = 4 G) Fraction of Homer1 puncta within CD68+ lysosomes in AAV-GFP and AAV-NPTX2 injected hippocampi of P301S mice. p value was determined by paired *t* test. n = 4 H) Relative spot number of Homer1 in AAV-GFP and AAV-NPTX2 injected hippocampi of P301S mice. p value was determined by paired *t* test. n = 4 Data are presented as mean ± SEM. Each dot represents data from individual samples.

### AAV-induced Nptx2 overexpression ameliorates synapse loss in P301S mice

Our data suggest that Nptx2 inhibits complement activation in the mouse brain and can limit neurotoxicity mediated by activated microglia. Therefore we considered whether overexpression of Nptx2 is sufficient to ameliorate synapse loss in a neurodegeneration mouse model characterized by glial activation, synapse loss and neurodegeneration. We showed previously in the TauP301S mouse model of FTD/AD (termed P301S mice hereafter) that C1q labels synapses for microglial elimination and that a C1q blocking antibody or genetic deletion of the downstream C3 is sufficient to ameliorate synapse loss (Dejanovic et al., 2018; Wu et al., 2019). To determine if overexpression of Nptx2 prevents synapse loss in P301S mice, we stereotactically injected AAVs overexpressing V5-tagged Nptx2 or GFP as a control into the ipsilateral and contralateral hippocampal hemisphere, respectively (Fig. 5A; S6A). We injected 8.5 months old P301S mice, an age when gliosis is present, complement is overactivated and synapses loss is detectable (Dejanovic et al., 2018), and analyzed the brains 6 weeks post-injection by IHC (Fig. 5B). As expected at this age, P301S brains were characterized by robust phospho-Tau staining (measured by AT8 immunoreactivity), elevated C1q immunoreactivity as well as microgliosis and astrogliosis as measured by Iba1 and GFAP immunoreactivity, respectively (Fig. S6B-F); all those hallmarks were most strongly detectable in the hippocampus as previously described (Dejanovic et al., 2018; Wu et al., 2019) (Fig. S6B). Nptx2 overexpression did not affect AT8, C1q, Iba1 and GFAP immunoreactivity (Fig. S6G-J). However, overexpression of Nptx2 reduced the percentage of C3-labeled synapses, suggesting that CCP activity is curbed as a result of increased Nptx2 expression (Fig. 5C, D). Consistent with restrained CCP activity, the volume of CD68+ microglial lysosomes was significantly smaller (Fig. 5E, F) and microglial engulfment of excitatory synapse was significantly reduced (Fig. 5E, G) in Nptx2 vs. GFP overexpressing hippocampal regions. This correlated with an increase in synapse density in Nptx2-overexpressing hippocampi, as measured by number of Homer1 puncta (Fig. 5H). Thus, Nptx2 overexpression is sufficient to ameliorate CCP-mediated microglial synapse pruning in P301S mice without affecting tau pathology or gliosis, implying that Nptx2 and complement act downstream of Tau pathology and microglial activation.

### Nptx2, C1q and Nptx2-C1q complex levels are altered in genetic FTD

Complement is recognized as a major driver of synapse loss and neuronal damage in many neurodegenerative diseases, and we reported previously that total and cleaved, i.e. activated, C3 is increased in CSF from AD patients (Wu et al., 2019). In a previously characterized genetic FTD patient cohort, symptomatic carriers of pathogenic mutations in *GRN* or *C9orf72* had significantly lower Nptx2 concentrations in CSF, as well as decreased Nptx1 and Nptxr levels, compared to presymptomatic carriers or non-carrier controls (Ende et al., 2020) (Fig. 6A). In a subset of this cohort, we measured CSF C1q concentrations by ELISA (Fig. 6B). While C1q concentrations were similar in CSF samples from non-carriers and presymptomatic carriers, they were significantly elevated on average in CSF from symptomatic patients (Fig. 6B). Next, we wanted to determine if C1q and Nptx2 present in the CSF are biochemically associated and if Nptx2-C1q complex levels are changed in FTD CSF. We utilized the PLA that we previously used to measure Nptx2-C1q complex levels in culture medium. The specificity in complex samples was validated using Nptx2^KO^ and C1q^KO^ mouse brain lysates (Fig. S7A). Furthermore, we monitored whether Nptx2 and C1q would form new protein complexes in solution by co-incubating Nptx2^KO^ and C1q^KO^ lysates. However, we detected only background signal compared to lysates from WT brains (Fig. S7A), suggesting that we primarily detect Nptx2-C1q complexes that pre-existed *in vivo*. Additionally, we confirmed that the assay requires the presence of both oligo-labeled C1q and Nptx2 antibodies in WT brain lysates (Fig. S7B). Lastly, compared to normal human serum, we measured only background Nptx2-C1q PLA signal in C1q-depleted human serum (Fig. S7C). Interestingly, using the C1q-Nptx2 PLA assay, we measured robust signal in the CSF, suggesting that C1q and Nptx2 are present in a complex in the CSF (Fig. 6C). The Nptx2-C1q PLA signal, i.e. the level of the protein-complex containing these two proteins, was significantly decreased in symptomatic vs presymptomatic and non-carrier CSF (Fig. 6C). To test if downstream complement activity is changed in FTD CSF, we measured activated C3b levels by ELISA. CSF C3b concentration strongly correlated with C1q levels (Fig. S7D). Importantly, compared to presymptomatic and non-carriers, C3b concentration was significantly increased in symptomatic CSF (Fig. 6D). By comparison, CSF levels of Factor B, a component of the alternative complement pathway, were similar between the three patient groups (Fig. 6E).

**Figure 6:**
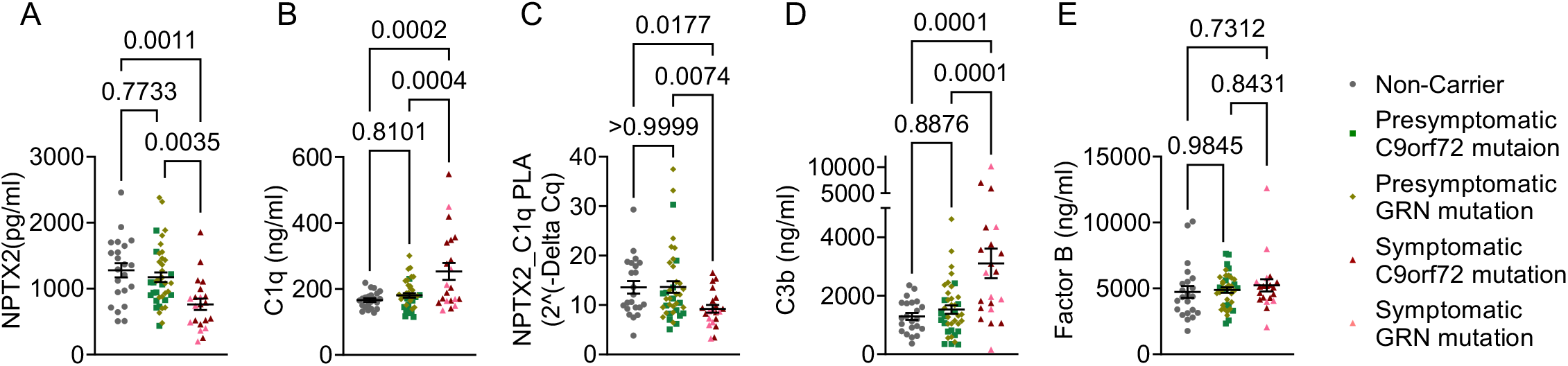
Nptx2, C1q and Nptx2-C1q complex levels are altered in genetic FTD. A, B, D, E) Concentrations of Nptx2, C1q, C3b and Factor B in cerebrospinal fluid (CSF) from non-carriers, presymptomatic and symptomatic carriers of *C9orf72* or *GRN* mutations. Protein levels were measured by ELISA. Nptx2 concentrations were measured and reported previously (Ende et al., 2020). P values were determined by one way ANOVA test. C) NPTX2-C1q complex levels in CSF samples analyzed by PLA. Quantitative qPCR levels were represented using 2^ (-delta Cq). See Methods for details. P values were determined by one way ANOVA test. All data are presented as mean ± SEM. Each dot represents data from one individual. Non-carriers, n = 22; presymptomatic GRN mutation, n = 25; presymptomatic C9orf72, n = 15; Symptomatic GRN mutation, n = 7; symptomatic C9orf72, n = 14.

Thus, in CSF from symptomatic genetic FTD patients Nptx2 concentration and Nptx2-C1q complex levels are lowered, while CCP activity is increased. These findings support that the role of Nptx2 as a potential regulator of CCP activity in the brain.

## Discussion

We report that Nptx2 binds C1q and thereby inhibits the CCP activity in the brain under physiological and pathological conditions. The CCP is triggered by binding of C1q to a wide range of receptors and subsequent activation of the downstream pathway. While other neuronal CCP inhibitors have been identified to play a role in synapse sculping during neurodevelopment, including SRPX2 and CSMD1 (Baum et al., 2020; Cong et al., 2020), to our knowledge Nptx2 is the first neuronal protein that regulates CCP activity in the adult brain. Deletion of Nptx2, which is reduced in CSF from schizophrenia, AD and FTD patients, increased CCP activity resulting in microglia-mediated removal of excitatory synapses in the adult brain. Consistent with the hypothesis that Nptx2 acts as a CCP inhibitor by functioning as a C1q decoy receptor, overexpression of Nptx2 was sufficient to ameliorate neural damage in a neuroinflammation cell model and in P301S mice. One important finding of our study is the diminished levels of Nptx2-C1q complexes and elevated C3b levels in the CSF from symptomatic genetic FTD subjects, which is indicative of elevated CCP activity in the brain at a stage of disease when cognitive failure is present.

A recent study reported that Nptx2 interacts with C1q’s globular head domain with a binding affinity in the nanomolar range (Vadászi et al., 2022), which independently validates our protein interaction analysis. Since Nptx2 binds to the head region of C1q, the domain by which C1q interacts with targets like antigen-bound synapses or IgGs, synaptic Nptx2 likely sequesters C1q and thereby blocks downstream CCP activation. A similar mode of action has been described for a therapeutical C1q-neutralizing antibody that binds to the head domain of C1q and prevents the interaction of C1q with diverse ligands (Lansita et al., 2017; Vukojicic et al., 2019), including synapses (Dejanovic et al., 2018) and is currently in clinical trials for HD and amyotrophic lateral sclerosis (ALS).

NPTXs bind postsynaptic AMPA receptors via a transsynaptic interaction, and, at least in neuronal cultures, this interaction appears to be sufficient to induce postsynaptic specializations (Lee et al., 2016; Xu et al., 2003). While we cannot exclude the possibility that Nptx2 might modulate synapse stability and/or density through binding to AMPA receptors or other (synaptic) interaction partners, the decreased synapse density in Nptx2^KO^ mice was C1q-dependent. The dynamic redistribution of synaptic Nptx2 in response to behavior, sleep, circadian rhythm, and afferent activity (Severin et al., 2021; Xiao et al., 2021) indicates that Nptx2-C1q might contribute to the precise refinement of circuits that mediate homeostasis of excitability, rhythmicity, and capacity for information processing.

Without a ‘stressor’ and subsequent glial cell activation and expression of downstream complement factors, C1q accumulation alone is likely insufficient to trigger aberrant innate immune reaction and widespread neuronal damage (Stephan et al., 2013; Wu et al., 2019). Our data suggest that in the absence of Nptx2, basal CCP activity is elevated enough to increase microglia-mediated excitatory synapse elimination without inducing more severe neuronal damage. This is consistent with previous work showing that mice deficient in the complement inhibitors SRPX2 or CSMD1 are characterized by loss of excitatory synapses without frank neurodegeneration early in brain development (Baum et al., 2020; Cong et al., 2020). Genetic variations of the complement component 4 (C4) gene *C4A*, which lead to increased expression of the C4a protein, is among the strongest common variant risk factors associated with schizophrenia (Consortium et al., 2016; Kamitaki et al., 2020). Increased expression of C4 is sufficient to elevate microglia-mediated elimination of excitatory synapses resulting in aberrant behavior in mouse models (Comer et al., 2020; Yilmaz et al., 2021). Nptx2^KO^ mice parallel the phenotype of C4-overexpressing mice; we found elevated levels of cleaved C4b in brains from Nptx2^KO^ mice and we previously reported that Nptx2^KO^ mice exhibit increased vulnerability to social isolation stress with emergent neuropsychiatric behavioral deficits (Xiao et al., 2021). The present finding extends the notion that Nptx2 loss of function spans the complex genetics of schizophrenia as a shared common pathway (Xiao et al., 2021).

In contrast to modestly raised CCP activity and moderate loss of excitatory synapses in Nptx2^KO^ or C4-overexpressing mice, C1q and downstream complement components are highly upregulated in many chronic neurodegenerative diseases, including AD (Dejanovic et al., 2018; Hong et al., 2016a; Wu et al., 2019), FTD (Lui et al., 2016), glaucoma (Stevens et al., 2007) and HD (Wilton et al., 2021). The marked increase in CCP activity in these diseases correlates with exacerbated synapse loss and severe neurodegeneration, and genetic or pharmacological inhibition of CCP factors is sufficient to ameliorate neuronal damage. Thus, the degree of CCP activity correlates with the severity of neuronal damage across multiple diseases with different etiologies and genetic risk factors. At least in our neuroinflammation neuron-microglia co-culture model, C1q activity is required for microglia-mediated neurodegeneration. In this context, it is remarkable that neuronal overexpression of Nptx2 in neuron-microglia cultures or P301S mice at an age when Tau pathology and neuroinflammation are present, limited complement-tagging and subsequent microglial removal of synapses. Our genetic FTD CSF analysis and recently published data from others (Ende et al., 2021, 2022) identifies that CCP activity is elevated in brains of symptomatic patients, potentially driven by the increased amount of C1q that is not controlled by Nptx2. The CSF results support the idea that, similar to our data in neuron-microglia co-cultures and P301S mice, Nptx2 binds C1q and thereby regulates CCP activity. Notably, CSF Nptx2 levels changes are detectable before CSF C1q and C3b concentrations increase at a later disease stage (Ende et al., 2021). Together with decreased C1q-Nptx2 complex levels that we measured in CSF from symptomatic *GRN* or *C9orf72* mutation carriers, the data supports the idea that decreased Nptx2 levels might unleash detrimental CCP activity. We hypothesize that beyond genetic FTD, CSF C1q-Nptx2 complex levels might be a disease-relevant biomarker across other CNS diseases characterized by aberrant CCP activity.

Overall, our study identifies Nptx2 as a neuronal synaptic CCP inhibitor. Elevating Nptx2 expression might be a viable therapeutic strategy to curb complement overactivation and subsequent neuronal damage in FTD, AD and other neuroinflammatory diseases.

## Acknowledgments

We thank the Janelia’s Virus Tool facility for helping prepare the AAV. We thank Richard Huganir’s lab (Johns Hopkins University) for kindly providing the PSD95-GFP^f/f^ mice. We thank Tzu-Ming Wang and David Hackos for help with creating the Nptxr/2 CHO cell lines. We also thank Ken Chan and Ben Deverman for helping with AAV design. We thank the members of the Worley’s laboratory, especially Dr. Wenchi Zhang and Liuqing Yang for insightful discussion of this project. We thank Dr. Raozhou Lin for insightful discussion of this project and valuable suggestions of the manuscript writing and editing. This study was supported by funding from the National Institutes of Health (Grant No. R35NS097966 and U19 AG065169 to P.F.W.).

## Material and methods

### Materials availability

All biological resources, antibodies, cell lines and model organisms and tools are either available through commercial sources or the corresponding authors. All unique/stable reagents generated in this study are available from the Lead Contact with a completed Materials Transfer Agreement. Further information and requests for resources and reagents listed in Key Resources Table should be directed to the Lead Contact.

### Animals

All animal maintenance and experimental procedures were conducted in accordance with to the policies and procedures described in the Guide for the Care and Use of Laboratory Animals of the National Institutes of Health and were approved by the Institutional Animal Care and Use Committees at the Broad Institute and at JHMI. Animals were group housed and maintained under standard, temperature-controlled laboratory conditions. All mice had free access to water and food and were housed with a 12h:12h light-dark cycle. Both male and female mice were used for all experiments. Nptx2^KO^ mice in congenic C57BL/6J background were obtained from Mark Perrin’s lab. PV-Cre mice (stock No: 017320), tdTomato reporter mice (stock No: 007914) and C1q^KO^ mice (stock No: 031675), TauP301S (stock No: 008169) were purchased from Jackson Laboratory. PSD95-GFP^f/f^ mice were generously provided by Dr. Richard Huganir’s lab at the JHMI.

### Cell culture and stable cell line

CHO cells were cultured in DMEM supplemented with 10% fetal bovine serum. CHO cells were incubated at 37 °C in 5% CO2. Stable cell line were generated as follows: A bicistronic expression vector was generated in the pTRE-tight-BI plasmid (Clontech) containing both human Nptx2 and human Nptxr cDNAs. This vector only allows expression of Nptx2 and Nptxr in the presence of doxycycline. This Nptx2-Nptxr expression vector and linear Hygromycin marker (Clontech) were cotransfected into CHO-tetON cells (Clontech). Stably expressing cell lines were selected for by the addition of 1mg/mL hygromycin B to the media and single colonies were picked to generate monoclonal stable cell lines. Cell lines were screened using RT-PCR for Nptx2 and Nptxr in the presence and absence of 5ug/mL doxycycline and the cell line showing the strongest doxycycline-dependent expression was selected. The final cell line was passaged using media containing 300 ug/mL G418 and 500 ug/mL hygromycin and induced with 5 ug/mL doxycycline. CHO cells were incubated with purified C1q (Complement Technology) and bound C1q was visualized by immunofluorescence labeling.

### AAV and Plasmids

The sequence of CamKII promoter was obtained from (Holehonnur et al., 2015). The sequence of miR-204, which we found to reduce AAV transgene expression in mice (unpublished data), was obtained from (Jovičić et al., 2013). Plasmid pAAV-CamKII-Nptx2-V5 was generated by GenScript, by cloning a synthesized CamKII-Nptx2-V5-miR204 DNA fragment upstream of the WPRE-hGHpA sequence of a plasmid derived from plasmid CAG-NLS-GFP (a gift from Viviana Gradinaru, Addgene #104061). Plasmid pAAV-CamKII-NLS-GFP was generated by replacing the Nptx2-V5 fragment between the KpnI and EcoRI sites of plasmid pAAV-CamKII-Nptx2-V5 with an NLS-GFP fragment excised from plasmid CAG-NLS-GFP. Plasmid sequences were verified by Sanger sequencing. Recombinant AAVs were produced and tittered as previously described (Krolak et al., 2022). Briefly, HEK 293T/17 cells (ATCC, CRL-11268) were cultured in Dulbecco’s Modified Eagle Medium with high glucose, sodium pyruvate, GlutaMAX, and Phenol Red (DMEM, Gibco 10569044) supplemented with 5% Fetal Bovine Serum (FBS, Gibco 16000044) and 1x MEM Non-Essential Amino Acids Solution (NEAA, Gibco 11140076), and triple transfected with plasmid DNA encoding rep-cap, pHelper, and an ITR-flanked transgene using polyethylenimine (PEI, Polyscience, 24765-1). AAVs were harvested from the cells and the media 3 days post transfection and purified by ultracentrifugation over iodixanol gradients. The concentration of packaged virus genomes was tittered by digital droplet PCR (ddPCR) using ITR primers (forward: 5’-GGAACCCCTAGTGATGGAGTT-3’, reverse: 5’- CGGCCTCAGTGAGCGA-3’, synthesized by IDT) and probe (5’-FAM-CACTCCCTC-ZEN-TCTGCGCGCTCG-IBFQ- 3’, synthesized by IDT).

#### AAVs used in vivo

pAAV-CaMKII-GFP (Addgene:64545) were purchased from Addgene. pAAV-CaMKII-NPTX2-V5 were generated using NEBuilder HiFi DNA assembly cloning kit (Cat.E5520S). Sequence was confirmed by sanger sequence. AAVs were prepared by the Janelia Viral Tools facility.

### Plate-based binding and complement activation assays

*Pentraxins Binding C1q* Purified recombinant neuronal pentraxin proteins (4 µg/mL, R&D Systems) or IgM (2 µg/mL, Thermofisher Scientific) were suspended in carbonate buffer (10 mM NaHCO_3_; pH 9.6) and added to 96-well Nunc MaxiSorp Flat-Bottom plates, which were sealed and left to incubate with shaking overnight at 4°C. After immobilization, the plate was washed with 0.05% Tween-20 in PBS (PBST) and blocked with 2% w/v ELISA-Grade BSA in PBS (2% BSA) with shaking for 1 hour at 37°C. The plate was washed with PBST, and purified human C1q protein (2 µg/mL, Complement Technology) was suspended in GVB++ Buffer (Complement Technology) and was added to plate with shaking for 1 hour at 37°C. The plate was washed with PBST and incubated with primary antibody against C1q (ThermoFisher Scientific) in 2% BSA for 1 hour with shaking at 37°C. The plate was washed with PBST and incubated with species-specific HRP-conjugated secondary antibody in 2% BSA with shaking for 1 hour at 37°C. The plate was washed with PBST and chemically exposed by adding 100ul TMB substrate (abcam). The reaction was stopped with 100ul Stop Solution 450 nm (abcam) and the OD_450_ was recorded using EnVision 2104 multilabel plate reader (Perkins Elmer).

*Pentraxins Binding to Complement Proteins* Same protocol used above except the following modifications. Purified human complement proteins (4 µg/mL, Complement Technology) were immobilized and were treated with His-tagged recombinant neuronal pentraxin proteins (4 µg/mL, R&D Systems). The readout was HRP-conjugated anti-His tag antibody recognizing the pentraxin proteins (abcam).

*Pentraxins Inhibiting NHS CCP* Same protocol used above except the following modifications. Purified IgM (4 µg/mL) or BSA was immobilized on the plate. Before addition to the immobilized IgM/BSA, normal human sera (Complement Technology) was incubated with different concentrations of purified recombinant human pentraxin protein (Nptxr) suspended in GVB++ Buffer. The mixture was incubated with shaking for 15 min at 37°C and then added to the immobilized IgM/BSA. The control was heat-inactivated NHS and C1q-depleted sera. The readout was a primary antibody against cleaved (active) C3 (Hycult Biotech) and a species specific HRP-conjugated secondary.

*Validation of C1q Blocking Antibody* Same protocol used above except the following modifications. Purified Nptx2 protein (4 µg/mL) or BSA was immobilized on the plate. Prior to addition to the plate, mouse sera (Complement Technology) was incubated with C1q (4.8) antibody (Abcam) or rabbit isotype control antibody (CST) suspended in GVB++ buffer and incubated with shaking for 30 min at 37°C. The readout was a primary antibody against mouse cleaved C3 (Hycult Biotech) and a species specific HRP-conjugated secondary.

### Protein purification and *in vitro* pull down

Rat Nptx2 pentraxin domain (X2-PD) was synthesized and codon optimized for expression in *E. coli*. The construct was cloned into the PEG30a-GB1-His and transfected into Shuffle T7 expression competent *E. coli* (NEB). The transfected *E. coli* cell culture was grown at 30 °C until OD600 nm reached 0.6. The temperature was reduced to 20°C, and cells were induced with 0.4 mM isopropyl-β-D-thiogalactopyranoside (IPTG) and grown for 16 h. The cells were harvested and lysed by sonicating in 10mM HEPES (pH 7.3) containing 150mM Nacl, supplemented with 1% Triton X-100. The lysate was clarified by centrifugation at 8000 × *g* for 30 min at 4 °C. The supernatant was filtered and loaded onto a 5ml HisTrap column (GE Healthcare). The column was washed with 10 column volumes of Buffer A (10 mM HEPES, pH 7.3, 0.15 M NaCl, 5 mM imidazole) on an Aktapurifier (GE Healthcare) while collecting fractions. To elute bound protein, a linear gradient from 0 to 100% Buffer B (10 mM HEPES, pH 7.3, 0.15 M NaCl, 500 mM imidazole) was set over 25 column volumes and protein was fractionated. Fractions were analyzed by SDS-PAGE, and which containing the interested protein were pooled and dialyzed against 2 liters of 10 mM HEPES, pH 7.3, 0.15 M NaCl, 5mM DTT at 4 °C overnight. The following day, the dialyzed proteins were concentrated and loaded onto a Superdex 200 size exclusion column. Fractions were analyzed by SDS-PAGE, and fractions containing pure X2-PD protein were pooled and concentrated. Rat Nptxr pentraxin domain (XR-PD) was synthesized and codon optimized for expression in *E. coli*. The construct was cloned into the PGEX-6P-1 vector and transfected into Shuffle T7 expression competent *E. coli* from NEB. The *E. coli* growth and induced protein expression was performed in the same manner with X2-PD. The recombinant protein in the supernatant of the cell lysate was mixed with Glutathione Sepharose resin (GE Healthcare) and the mixture was washed extensively in 10mM HEPES pH 7.3. GST-XR-PD was eluted by 10mM GSH and loaded onto a Superdex 75 size exclusion column. Fractions were analyzed by SDS-PAGE, and fractions containing pure XR-PD protein were pooled and concentrated.

### *E. coli* killing assay

*E. coli* cells used in these assays were obtained from cultures grown in LB medium. Different amounts of XR-PD (0.5-5 µM), together with 0.25% NHS or inactivated NHS were preincubated for 10 min at 37 °C before adding *E. coli*. After an incubation of 30 min at 37 °C cells were plated to a LB-agar plate and cultivated overnight at 37 °C before counting colony forming units. Colony forming unit in GVB++ buffer was set as 100%.

### Hemolytic assay

To analyze inhibition of hemolysis via the classical pathway, 0.5% normal human serum (NHS), various concentrations of GST or GST-XR-PD were preincubated together for 15 min at 37 °C. After preincubation, the NHS–protein mix was added to the erythrocytes and incubated for additional 30 min at 37 °C. Lysis of erythrocytes was determined by measuring the amount of hemoglobin in the supernatants at 414 nm. Erythrocytes (EA) treated with ddH_2_O or incubated with GVB++ buffer served as positive (100% Hemolytic) and negative controls (0% Hemolytic).

### Western blot

Brain tissue and cultured cells were lysed in RIPA buffer. Protein samples were boiled in 1x reducing agent and 1x sample buffer (Invitrogen) and separated by 4-12% SDS-PAGE (ThermoFisher Scientific). Gels were transferred to PVDF membranes (ThermoFisher Scientific) and blocked with 5% milk in 0.1% Tween 20 (v/v) in TBST buffer with rocking for 1 hour at room temperature (RT). Primary antibodies in 1% Milk (w/v) in TBST were incubated with rocking overnight at 4°C. The membrane was washed with TBST and incubated with secondary HRP-conjugated antibodies in 1% Milk with rocking for 1 hour at RT. The membrane was washed, exposed with West Pico ECL (ThermoFisher Scientific), and immunoreactivity was detected on ChemiDoc system (Bio-Rad). Analysis was performed using Image Lab software (Bio-Rad).

### Electron microscopy negative staining

The single C1q protein particles, single Nptx2 protein particles, and C1q-NPTX2 complex was visualized by negative staining and electron microscopy followed the published protocol as described (Zhang et al., 2013). Briefly, to visualize C1q and Nptx2 single particles, purified C1q and Nptx2 were diluted to 5 μg/ml in 50 mM HEPES containing 150 mM NaCl; to visualize complex C1q-NPTX2, two proteins was directly incubated at 4°C for 2 hours and diluted to 5 μg/ml in 50 mM HEPES containing 150 mM NaCl. Carbon-coated grids were hydrophilized by glow discharge at low pressure in air. Aliquots of C1q, Nptx2, and C1q– Nptx2 complex were adsorbed onto hydrophilic, carbon-coated grids for 1 min, washed twice with ddH2O, and stained on a drop of 2% Uranyl Formate (UF) in ddH_2_O. Specimens were examined in a Hitachi 7600 TEM at a 300,000x magnification.

### ELISA assay

#### Mouse brain lysates

Frozen brain tissue was hand homogenized in lysis buffer (5mM HEPES pH 7.4, 1mM MgCl2, 0.5mM CaCl2, supplemented with protease inhibitors (cOmplete protease inhibitor cocktail, Roche). After low speed centrifugation (1,400 xg, 10 min, 4C), the post-nuclear supernatant fractions were used to measure mouse C1q, C4 and C4b levels using commercial ELISA kits; C1q mouse ELISA (Hycult Biotech, HK211-01), Total C4 (LSBio, LS-F37428-1), and C4b (LSBio, LS-F8056-1) following the manufacturer’s instructions. Brain lysate protein concentration was measured using a BCA kit (Thermo Scientific). The experimenters were blinded to the genotype.

#### Human CSF samples

CSF complement proteins C1q, C3b and Factor B were measured using the ELISA kits Human complement C1q (ab170246), Human Complement C3b ELISA kit (ab195461), Human Complement Factor B ELISA Kit (ab137973) from Abcam according to the manufacturer’s instructions. All the CSF measurements were performed in duplicate and blinded to all clinical and genetic information.

### Antibodies and AAVs injection

C1q (4.8) antibody and isotype control antibody injection were performed as described previously with slight modification (Dejanovic et al., 2018). Briefly, the C1q (4.8) antibody and Rabbit IgG isotype control (Invitrogen: Cat 10500C) were concentrated to 6.7 mg/ml. 2-3 months old Nptx2^KO^ mice were anesthetized with 3% isoflurane and placed on a stereotaxic apparatus (RWD) for surgery. All injections were performed with Nanoject II (Drummond scientific company). The injection coordinate is M/L=+/-3.2 mm, A/P=0.02mm, D/V= 2.5mm. Two μl of anti-C1q or isotype control antibody were injected at a rate of 50.6nl/per injection. The needle was removed 5 minutes after completion of the injection and mice were put back in their cage. Six days after injection, the mice brains were harvested, and IHC staining, imaging and data analysis were performed.

For AAV virus injection of TauP301S mice, we performed bilateral injection with the AAV-CamkII-GFP as control and AAV-CamkII-NPTX2-V5 in both cortex and Hippocampus. The right hemisphere was injected with AAV-CamkII-GFP and the coordinate is M/L=-2.5 mm, A/P=-2.18mm, D/V= -2.2mm; the left hemisphere was injected with AAV-CamkII-NPTX2-V5 and the coordinate is M/L=2.5 mm, A/P=-2.18mm, D/V= -2.2mm; As the coordinate for cortex injection is M/L=+/-2.5 mm, A/P=-2.18mm, D/V= -1.1mm. Mice brains were harvested 6 weeks after injection and IHC staining, imaging and data analysis were performed.

### Immunohistochemistry staining

Mice brains were harvested following transcranial perfusion with PBS and 4% paraformaldehyde (PFA). Tissue was postfixed in 4% PFA overnight, then washed with PBS and transferred to 30% sucrose solution until the tissue has sunk to the bottom of the tubes. Post-fixed brains were sliced in 35 μm thick sections in a -18°C chamber using a 3050 S Cryostat (Leica). Brain slices were either placed directly on Superfrost Plus slides (VWR) and stored at -80°C until ready for use or directly put in the PBS and store in the cold room. Mice brains slice were blocked with blocking buffer (either 10% Normal goat serum, 1% BSA, 0.3% tween-20 in PBS or 5% BSA, 0.2% Triton X-100 in PBS) for 1 hour, and incubated with primary antibodies overnight. After rinsing with PBS four times, secondary antibodies were applied and incubated for 1 hours at room temperature. Slides were then washed and mounted using the ProLong Gold Antifade Reagent with DAPI or without DAPI dependent on the specific samples.

### Microglial engulfment analysis

For analysis of microglial synapse engulfment, brain sections were imaged on the Zeiss LSM 880 confocal microscope or Andor DragonFly Spinning Disk Confocal Microscope with z-stacks (∼15 μm) using 63x oil objective. CD68-positive lysosomes were 3D-reconstructed using the surface rendering function in Imaris 9.7. Excitatory synapse (Vglut1 and Homer1) and Inhibitory synapse (Gephyrin and GAD67) puncta inside the CD68+ lysosomes were quantified.

### Synapse density analysis

Excitatory and inhibitory synapse density was analyzed using a custom script in Fiji (ImageJ) software followed the published protocol as described (Favuzzi et al., 2021). Briefly, tissue samples were imaged on a Zeiss LSM 880 confocal microscope or Andor DragonFly spinning disk confocal microscope using a 63x oil immersion objective. Images were processed and automatically analyzed using the script provided by Emilia Favuzzi, which can be found here: https://github.com/emiliafavuzzi/synaptic-analyses.

### Immunocytochemistry, imaging, and quantification in microglia-neuronal co-cultures

Rat neuronal cultures were prepared from hippocampi of rat embryos on embryonic day 18 (E18) or E19 and plated on poly-D-lysine and laminin coated coverslips at a density of 90,000/24-well dish and cultured in NbActiv4 medium (BrainBits). 50% of the medium was exchanged with fresh medium weekly. For primary microglial cultures, embryonic E18-E19 or postnatal (P1-P2) pups were decapitated, and forebrain was triturated with a 5 ml serological pipette and the homogenate was spun at 300xg for 5 min. The supernatant was discarded and the pellet was resuspended with a 1 ml pipette and filtered through a 75 μm filter. Two brains were cultured per 175 cm2 flask in 40 ml DMEM + 10% fetal bovine serum (FBS). After 24 hours incubation, flasks were rinsed with Hank’s Balanced Salt Solution (HBSS, Thermo Fisher Scientific) and new media was added. Cultures were grown for an additional 7-9 days before microglia were shaken off the astrocyte feeder layer on a rocking platform for 1 h, pelleted and resuspended in NbActiv4 medium and added to neurons in a 1:2 (microglia : neuron) ratio. Primary cells from male and female pups were used. 10k primary rat microglia were added to 20k DIV12 primary hippocampal neurons with 40 ng/mL of rat macrophage colony stimulating factor (MCSF) (Peprotech 400-28-100UG). At the time of plating, microglia were added to either 3-days post AAV-infected neurons (see AAV infection below) or DIV12 neurons in the presence of recombinant protein (see recombinant protein below) or C1q-blocking antibody (Recombinant Anti-C1q antibody (ab182451)) / IgG (Normal Rabbit IgG,Cell Signaling Technology). For LPS induced activation of microglia, LPS was added at a concentration of 1 ug/ml eBioscience™ Lipopolysaccharide (LPS) Solution (500X) (thermofisher)) 24 hours post seeding of microglia. 24-hours post treatment of LPS, media was collected, and cells were washed with 1X PBS and fixed with 4%PFA for imaging.

For AAV Infection, neurons were infected with AAV-GFP or AAV-NPTX2-V5 at DIV9 at a concentration of 100 viral genomes per cell (vg/cell). For recombinant NPTX2 treatment, DIV12 hippocampal neurons were treated with either 2 ug/mL of recombinant Nptx2 (CF 7816-NP-050, R&D Systems) or BSA as control. For antibody treatment, DIV12 hippocampal neurons were treated with either 1 ug/mL of C1q-blocking antibody or isotype control IgG.

Neuron-microglia co-cultures were fixed after blocking antibody addition, recombinant nptx2 or AAV expression and subjected to immunocytochemistry, using anti-MAP2 (1:500) (Abcam ab5392), anti-V5 (1:500) (MA5-15253, thermofisher), and anti-Iba1 (1:500) (Wako sc-133172). Slides were imaged with a Andor DragonFly Spinning Disk Confocal Microscope; Images were collected, processed using ImageJ, and quantified blind to conditions.

### Proximity ligase assay (PLA)

PLA was used as previously reported with some modification (Darmanis et al., 2010). Briefly, Nunc MaxiSorp 384-well plate (MilliporeSigma, cat# P6366-1CS) was coated with rabbit anti-Nptx2 (7 μg/ml, 25 ul/well) in carbonate-bicarbonate coating buffer (pH 9.6), and incubated overnight at 4°C. The next day, plates were washed twice with TBST and blocked with 5% BSA + 0.1 mg/ml salmon sperm DNA (ThermoFisher, cat# 15632011), 100ul/well, at room temperature (RT) for one hour. 20 μl of sample or 20ul TBS + 5% BSA + 0.1 mg/ml sperm DNA (as blank) was added to each well and incubated overnight at 4°C. After washing with TBST (5 times, 100 ul/well each time), 20 μl of PLA probe (1 μg/ml of oligo plus probe + 1 μg/ml oligo minus probe in TBS with 5% BSA + 0.1 mg/ml sperm DNA) was added, and incubated at RT for one hour. After washing with TBST (5 times, 2 min each, speed 1500 rpm on shaker), 20 μl of ligation mixture (0.4 μl of T4 ligase, 0.2 μl of connector oligo (20 μM), 2 μl of 10x T4 ligase buffer and 17.4 μl of H2O) was added. The plate was incubated at 37°C for 30 min, then at RT for one hour. After washing with PBS (3 times), add 20 μl of 20mM DTT and incubate at RT or 37°C for one hour. The eluted ligation products were collected and dilute with 20 ul H_2_O as qPCR template and qPCR was performed in 384-well plate (ThermoFisher cat# 4309849). The delta-delta Ct method was used for calculating PLA value in qPCR. Sequences of oligos

Oligo plus (59bp, HPLC purified, 5’ add amino-5AmMC6): 5’ CGCATCGCCCTTGGACTACGACTGACGAACCGCTTTGCCTGACTGATCGCTAAATCGTG 3’

Oligo minus (61bp, HPLC purified, 5’ add phosphorylation, 3’ add amino-3AmMO):

5’TCGTGTCTAAAGTCCGTTACCTTGATTCCCCTAACCCTCTTGAAAAATTCGGCATCGGTGA 3’

qPCR Primer 1 : 5’ CATCGCCCTTGGACTACGA 3’

qPCR Primer 2 : 5’ GGGAATCAAGGTAACGGACTTTAG 3’

Connector: 5’ TACTTAGACACGACACGATTTAGTTT 3’

### Statistical analysis

Number of samples is shown in figure and described in figure legends. All values are presented as mean ± SEM unless otherwise stated. Statistical analysis was performed using GraphPad Prism 9 and specific statistical tests are defined in figure legend. For all experiments, statistical significance is defined as follows: n.s. – not significant, *p < 0.05, **p < 0.01, ***p < 0.001.

## Figure legends

**Supplemental Figure 1.**
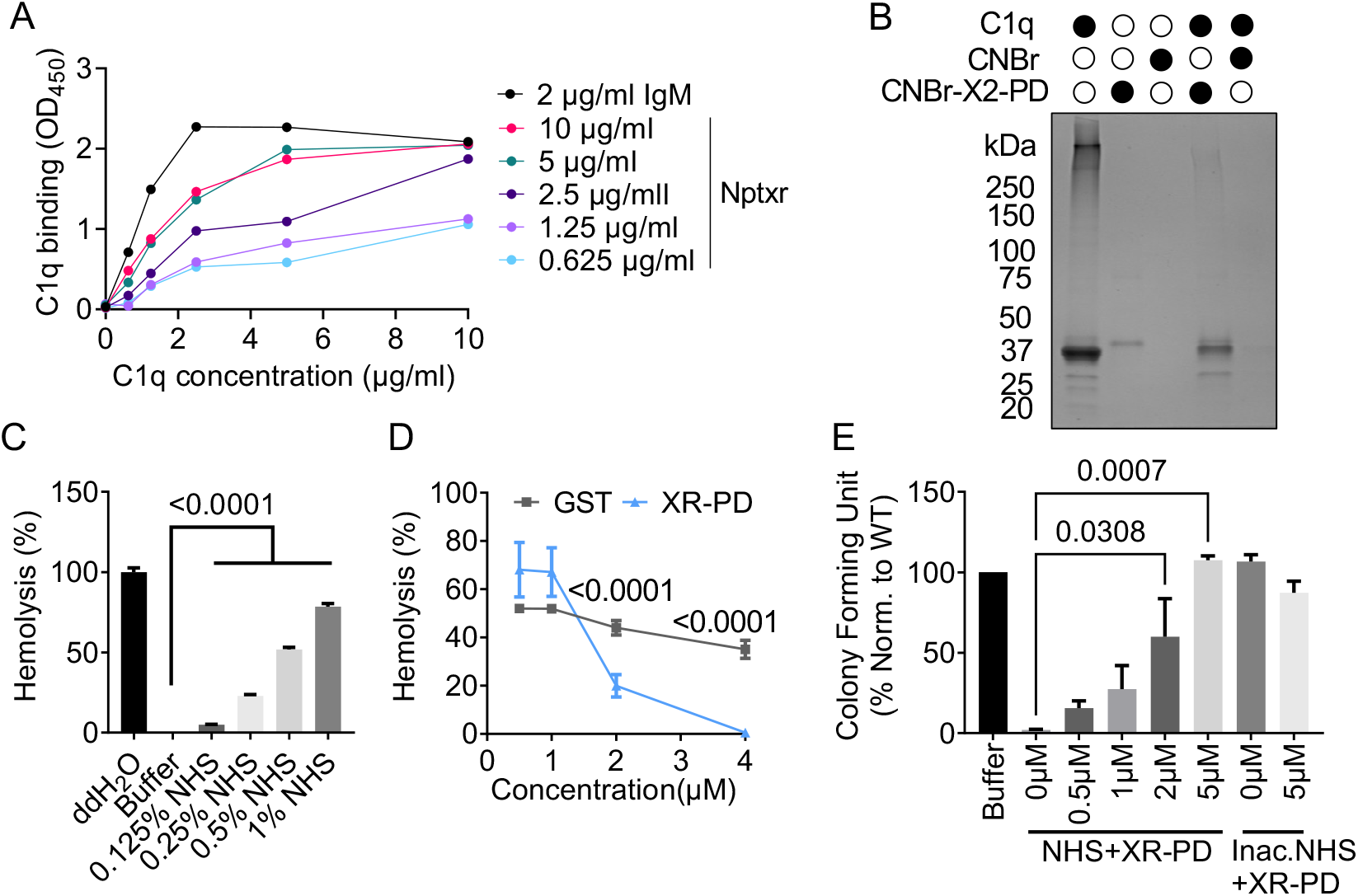
A) Microtiter wells were coated with 2µg/ml IgM or 0.625 – 10 µg/ml Nptxr and incubated with various concentrations of C1q as indicated. Bound C1q was detected using an anti-C1q antibody. B) Purified Nptx2 pentraxin domain (X2-PD) coupled to CNBr-activated Sepharose 4B was incubated with purified C1q. Proteins were separated by SDS-PAGE and visualized by Coomassie blue staining. C) Quantification of hemolysis of sheep erythrocytes in presence of increased NHS concentrations. Lysis of sheep erythrocytes was monitored by measuring released hemoglobin at 415 nm. p values were determined by one-way ANOVA. D) Purified XR-PD was added to NHS in a CCP-specific buffer. Lysis of sheep erythrocytes was monitored by measuring released hemoglobin at 415 nm. p values were determined by two-way ANOVA. E) Survival of *E. coli* in presence of purified XR-PD in NHS. *E. coli* survival was analyzed by counting colony forming units. Heat inactivated NHS was used as a negative control. p values were determined by one-way ANOVA. Data are presented as mean ± SEM. Each dot represents data from individual samples.

**Supplemental Figure 2.**
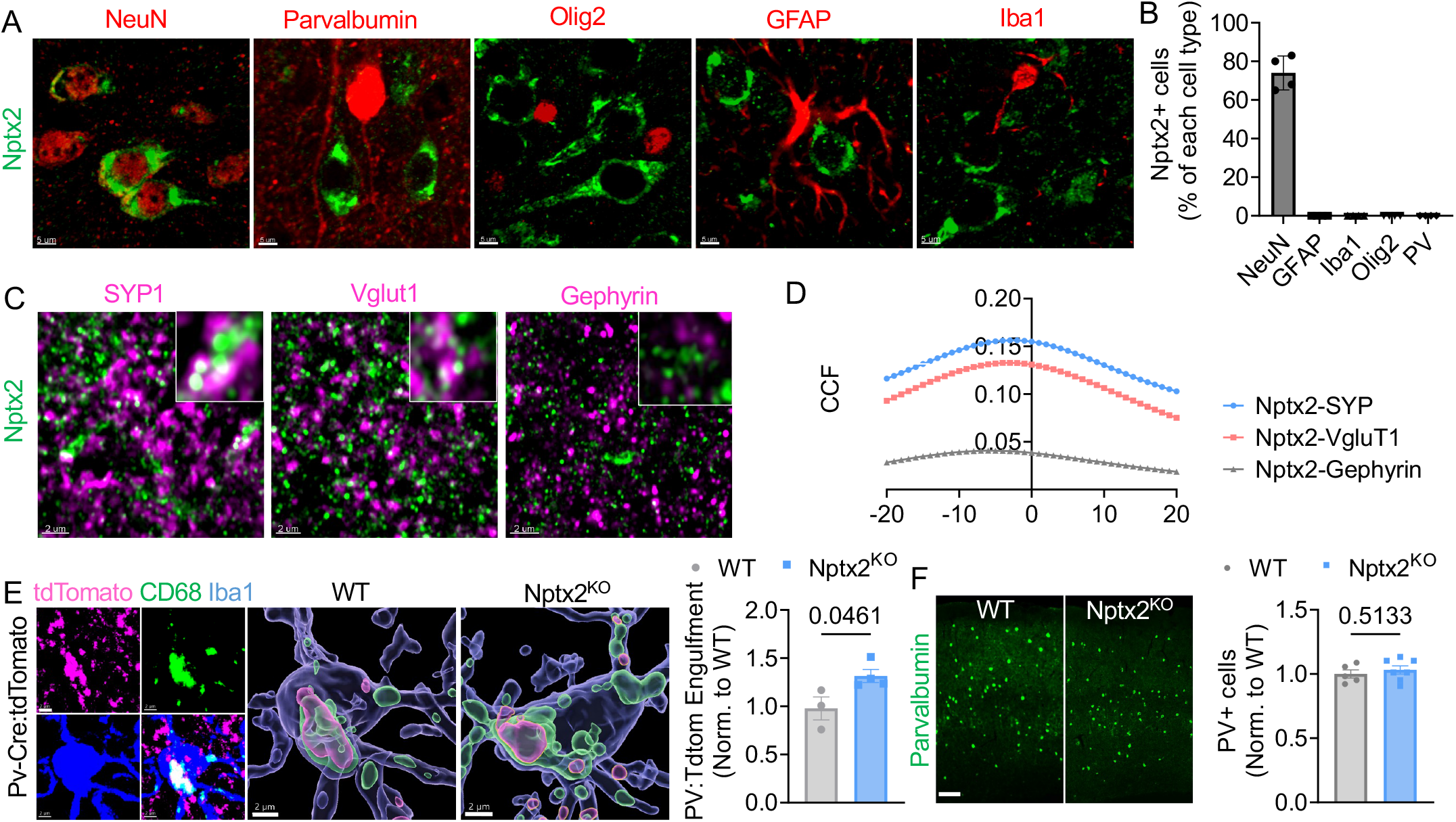
A) Representative confocal images of co-immunostained Nptx2 (green) with NeuN, Parvalbumin, Olig2, GFAP or Iba1 (all shown in red) in WT cortex. Scale bars are 5 µm. B) Percentage of Nptx2+ NeuN, Parvalbumnin, Olig2, GFAP and Iba1 cells in WT cortex. C) Representative confocal images of co-immunostained Nptx2 with the excitatory synapse marker proteins Synaptophysin 1 (SYP1) or Vglut1, and inhibitory synapse maker Gephyrin in WT cortex. Scale bars are 2 µm. D) Quantification of Nptx2 colocalization with SYP1, Vglut1 and Gephyrin. A Pearson’s coefficient cross-correlation function (CCF) was used to analyze the colocalization by calculated with the magenta channel shifted pixel by pixel with respect to red channel with ImageJ JACoP plugin. E) Representative image of tdTomato fluorescence and co-immunostained CD68+ microglial lysosomes (green) and Iba1 (blue). The three-dimensional surface rendering shows CD68+ microglial lysosomes containing tdTomato in PV-Cre:tdTomato mice in WT and Nptx2^KO^ background. Scale bars are 2 μm.Quantification of tdTomato volume within CD68+ microglial lysosomes in PV-Cre:tdTomato mice in WT and Nptx2^KO^ background. Values were normalized to WT mice. p value was determined by unpair *t* test. n = 3 for WT; n = 4 for Nptx2^KO^. F) Representative images and quantification of PV+ cell density in WT and Nptx2^KO^ cortex. Cell density was normalized to WT mice. p value was determined by unpaired *t* test. n = 5 for WT; n = 7 for Nptx2^KO^. Scale bar: 100µm. Data are presented as mean ± SEM. Each dot represents data from one mouse.

**Supplemental Figure 3.**
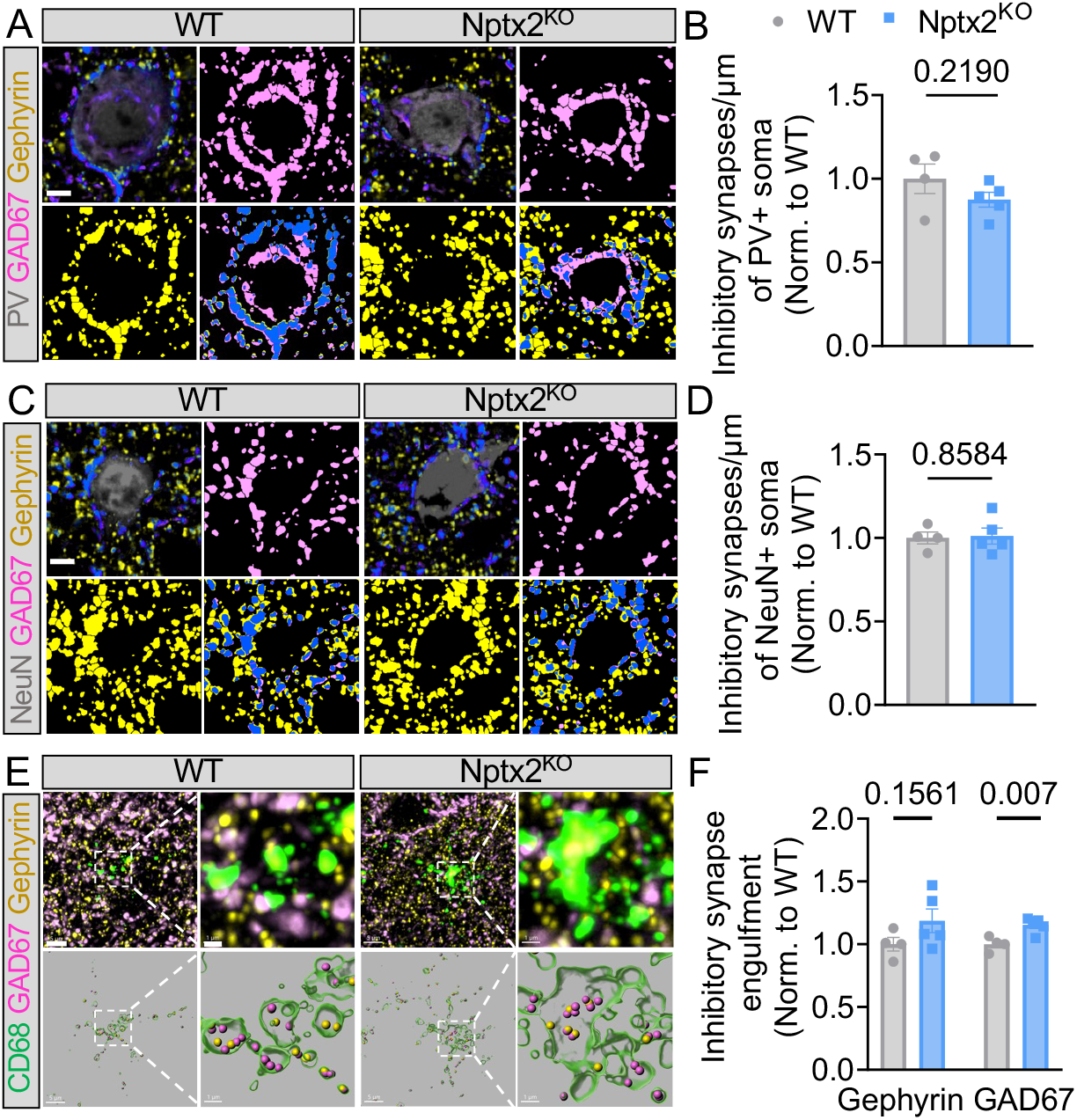
A, B) Representative confocal images of PV (gray), Gephyrin (yellow) and GAD67 (pink), and corresponding masks. Bar graph shows inhibitory synapse density around somas of PV+ neurons. p value was determined by unpair *t* test. n = 4 for WT; n = 5 for Nptx2^KO^. Scale bars is 5 µm. C, D) Representative confocal images of NeuN (gray), Gephyrin (yellow) and GAD67 (pink), and corresponding masks. Bar graph shows inhibitory synapse density around somas of NeuN+ neurons. p value was determined by unpair *t* test. n = 4 for WT; n = 5 for Nptx2^KO^. Scale bars is 5 µm. E, F) Representative confocal images and 3D rendering of CD68 (green), Gephyrin (yellow) and GAD67 (pink) in WT and Nptx2^KO^ cortex. Bar graph shows the relative amount of Gephyrin and GAD67 puncta within CD68+ microglial lysosomes. p values were determined by one-way ANOVA. n = 4 for WT; n = 5 for Nptx2^KO^. Scale bars are 5 µm, insets, 1 µm. Data are presented as mean ± SEM. Each dot represents data from one mouse.

**Supplemental Figure 4.**
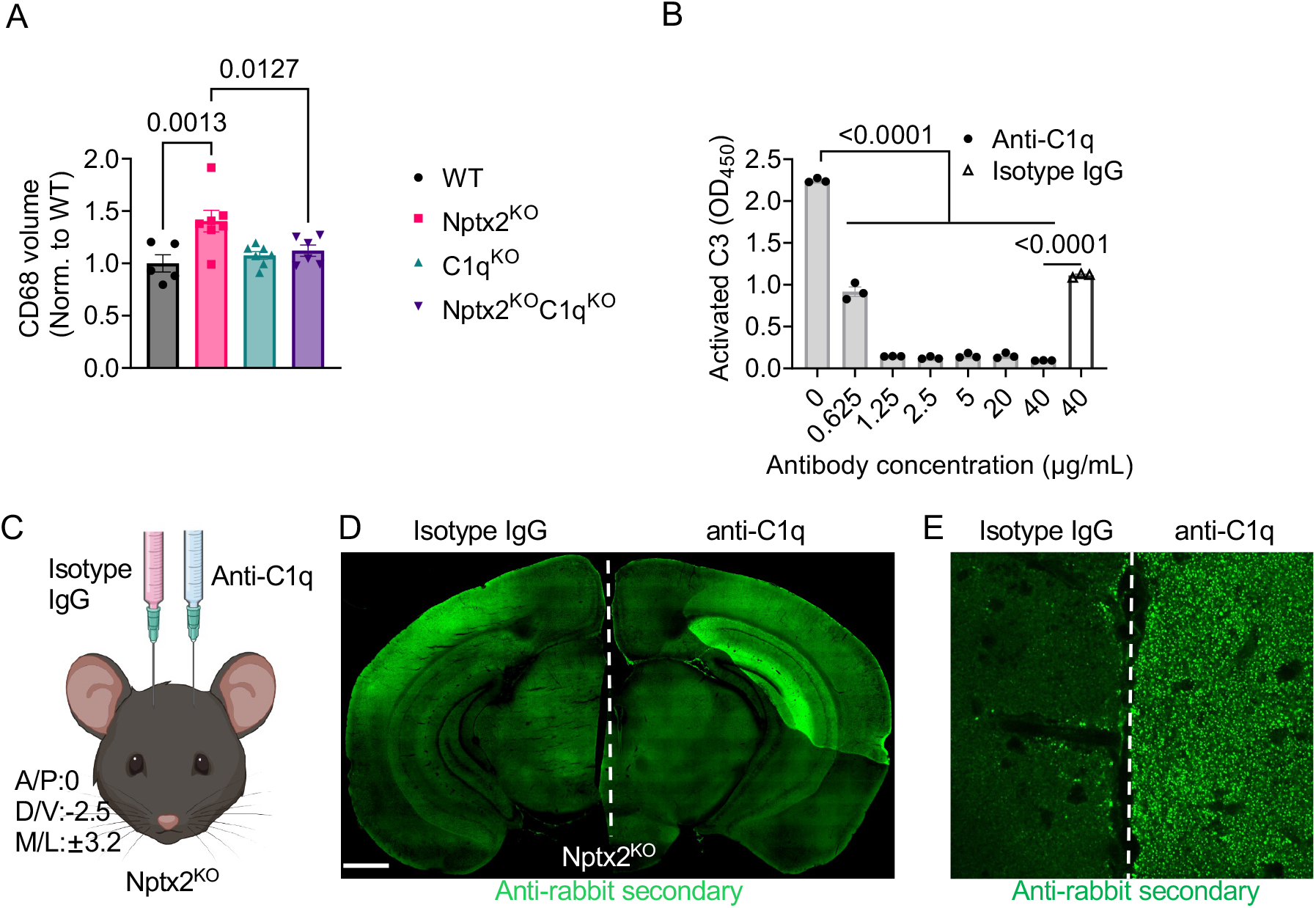
A) Volume of CD68+ microglial lysosomes in the cortex of mice with indicated genotypes. p values were determined by one-way ANOVA. n = 5 for WT; n = 7 for Nptx2^KO^; n = 7 for C1q^KO^; n = 6 for Nptx2^KO^;C1q^KO^. B) Nptx2-coated microtiter plates were incubated with 5% NHS supplemented with different concentrations of anti-C1q antibody. Complement activation and deposition was measured using an activated C3 specific antibody. Isotype IgG was used as negative control. p value were determined by one way ANOVA. C) Schematic of stereotactic antibody injection. Coordinates correspond to the injection site. D) Representative image of a Nptx2^KO^ brain injected with anti-C1q antibody and isotype IgG and immunostained with an Alexa-conjugated anti-rabbit secondary antibody. Scale bar is 1000 µm. E) Representative confocal image of a Nptx2^KO^ brain injected with anti-C1q and isotype IgG. Injected antibodies were detected with an Alexa-conjugated anti-rabbit secondary antibody. Note that while the immunoreactivity in the isotype IgG injected hemisphere is low and dim, the anti-C1q injected hemisphere shows bright puncta that is a characteristic C1q pattern. Data are presented as mean ± SEM. Each dot represents data from individual samples.

**Supplemental Figure 5.**
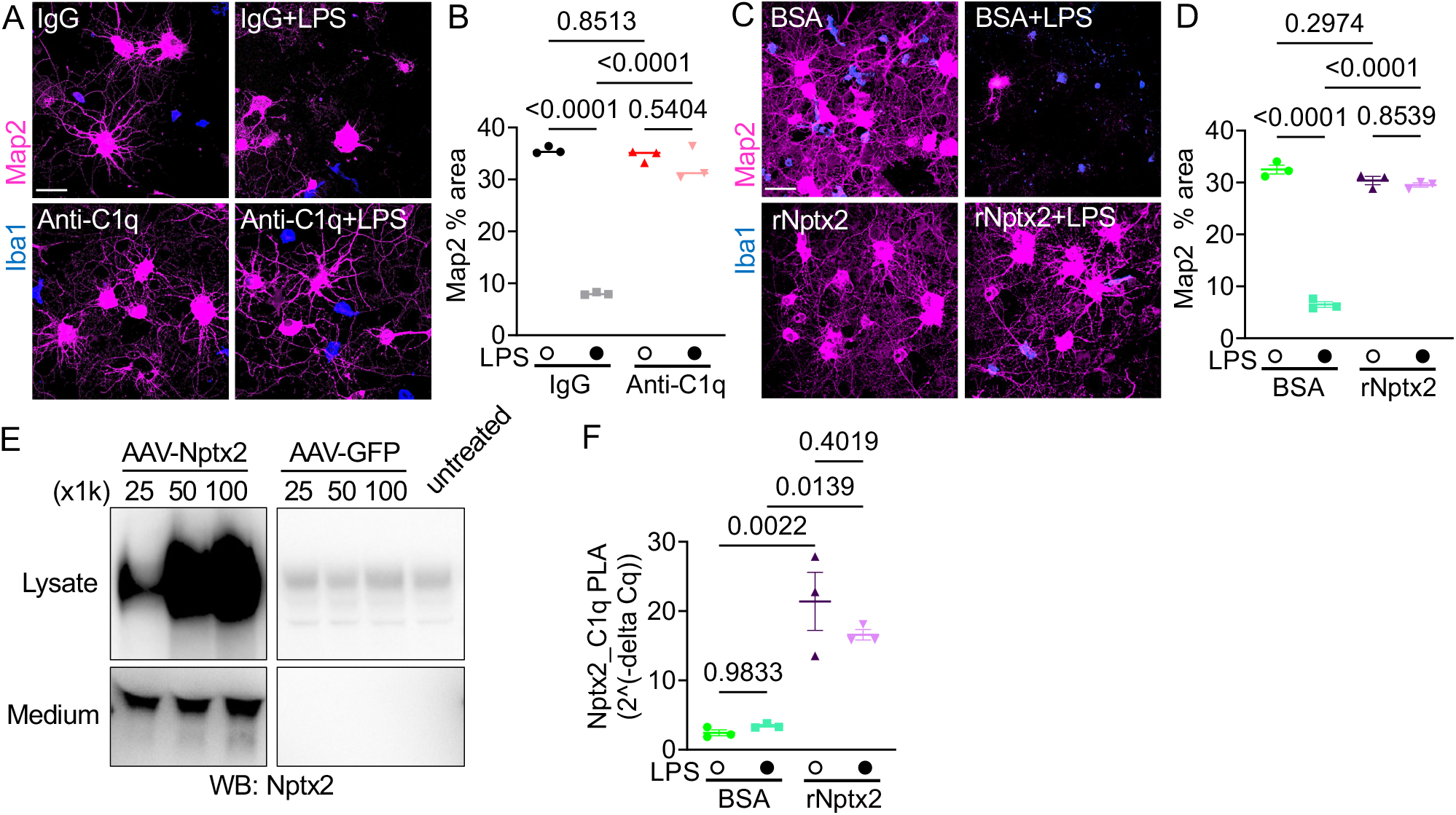
A) Representative images of immunostained Map2 (pink) and Iba1 (blue) in neuron-microglia co-cultures treated with LPS or vehicle control for 24 hours in the presence of control IgG or anti-C1q antibodies. Scale bar is 50 μm. B) Percentage of Map2+ area in indicated cultures. Three independent neuron-microglia co-cultures were used for the experiment. p values were determined by two-way ANOVA. C) Representative images of immunostained Map2 (pink) and Iba1 (blue) in neuron-microglia co-cultures treated with LPS or vehicle control in the presence of BSA or recombinant Nptx2 (rNptx2). Scale bar, 50 μm. D) Percentage of Map2+ area in indicated cultures. Three independent neuron-microglia co-cultures were used for the experiment. p values were determined by two-way ANOVA. E) Representative Western blot showing Nptx2 levels in lysate and culture medium of neurons infected with different virus titers (genome vector/per cell) of AAV-Nptx2 or AAV-GFP. F) Nptx2-C1q complex levels in culture medium analyzed by proximity ligation assay (PLA, see Methods). Media from three independent neuron-microglia co-cultures were used. p values were determined by two-way ANOVA.

**Supplemental Figure 6.**
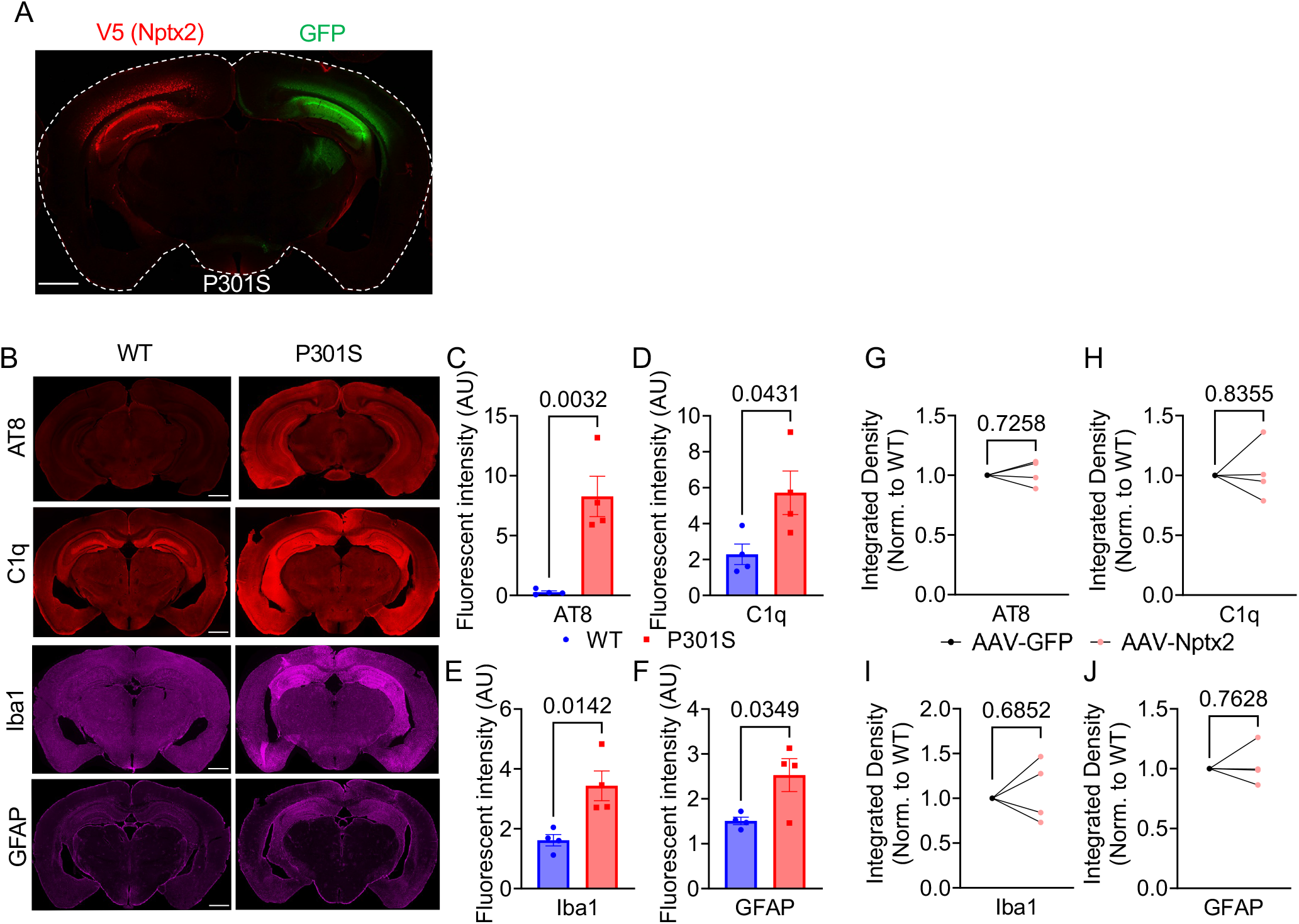
A) Representative confocal image of GFP and Nptx2 (detected by immunostaining the V5-tag) fluorescence in P301S brains injected with AAV-Nptx2 and AAV-GFP. Scale bar is 1000 μm. B) Representative images of WT and P301S brains immunostained with phospho-Tau (AT8), C1q, Iba1 and GFAP. Scale bar is 1000 μm. C-F) Immunofluorescence levels of indicated immunostainings in WT and P301S brains. p values were determined by unpaired *t* test. G-J) Immunofluorescence levels of indicated immunostainings in AAV-GFP and AAV-Nptx2 injected hippocampi of P301S mice. p values were determined by paired *t* test. Data are presented as mean ± SEM. Each dot represents data from individual samples.

**Supplemental Figure 7.**
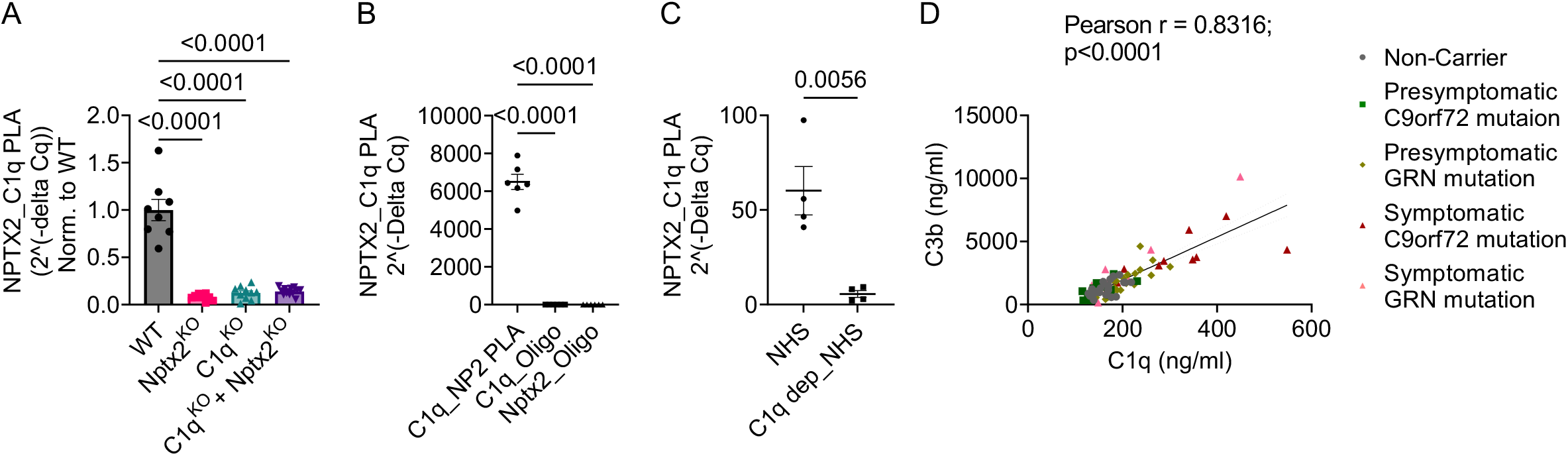
A) NPTX2-C1q complex levels in brain lysates from WT, Nptx2^KO^ and C1q^KO^ measured by PLA. Nptx2^KO^ + C1q^KO^ represent mixed Nptx2^KO^ and C1q^KO^ lysates. Quantitative qPCR levels are represented using 2^(-delta Cq). p values were determined by one way ANOVA test. n = 6 for WT, n = 6 for Nptx2^KO^. B) NPTX2-C1q complex levels in WT mouse brain lysates detected in presence of mixed oligo-labeled anti-C1q and anti-Nptx2 antibodies or in presence of individual anti-C1q or anti-Nptx2 antibodies. Quantitative qPCR levels are represented using 2^(-delta Cq). p values were determined by one way ANOVA test. n = 6 samples per condition. C) NPTX2-C1q complex levels in normal human serum (NHS) and C1q-depleted NHS. Quantitative qPCR levels are represented using 2^(-delta Cq). p value was determined by unpaired t test. n = 4 for NHS, n = 4 for C1q-depleted NHS. D) Correlation between C3b and C1q levels in CSF samples. P value was determined by two-tailed t test. Data are presented as mean ± SEM. Each dot represents data from one individual.

